# Connectomes across development reveal principles of brain maturation

**DOI:** 10.1101/2020.04.30.066209

**Authors:** Daniel Witvliet, Ben Mulcahy, James K. Mitchell, Yaron Meirovitch, Daniel R. Berger, Yuelong Wu, Yufang Liu, Wan Xian Koh, Rajeev Parvathala, Douglas Holmyard, Richard L. Schalek, Nir Shavit, Andrew D. Chisholm, Jeff W. Lichtman, Aravinthan D.T. Samuel, Mei Zhen

**Affiliations:** Lunenfeld-Tanenbaum Research Institute, Mount Sinai Hospital, Toronto, ON, Canada; Department of Molecular Genetics, University of Toronto, Toronto, ON, Canada; Department of Physics, Harvard University, Cambridge, MA; Center for Brain Science, Harvard University, Cambridge, MA; Computer Science and Artificial Intelligence Laboratory, Massachusetts Institute of Technology, MA; Division of Biological Sciences, Section of Cell and Developmental Biology, University of California, San Diego, CA; Department of Molecular and Cellular Biology, Harvard University, Cambridge, MA; Department of Physiology, University of Toronto, Toronto, ON, Canada

## Abstract

From birth to adulthood, an animal’s nervous system changes as its body grows and its behaviours mature. The form and extent of circuit remodelling across the connectome is unknown. We used serial-section electron microscopy to reconstruct the full brain of eight isogenic *C. elegans* individuals across postnatal stages to learn how it changes with age. The overall geometry of the brain is preserved from birth to adulthood. Upon this constant scaffold, substantial changes in chemical synaptic connectivity emerge. Comparing connectomes among individuals reveals substantial connectivity differences that make each brain partly unique. Comparing connectomes across maturation reveals consistent wiring changes between different neurons. These changes alter the strength of existing connections and create new connections. Collective changes in the network alter information processing. Over development, the central decision-making circuitry is maintained whereas sensory and motor pathways substantially remodel. With age, the brain progressively becomes more feedforward and discernibly modular. Developmental connectomics reveals principles that underlie brain maturation.

## Introduction

The developing nervous system faces multiple challenges. Amid an animal’s changing anatomy and fluctuating environment, some circuits must maintain robust outputs, such as locomotion^1–4^. New circuits need to be constructed in order to support new functions, such as reproduction^5–7^. To adapt and learn, the nervous system must make appropriate changes in existing circuits upon exposure to internal and external cues^8^.

The nervous system employs a variety of adaptive mechanisms to meet these challenges. In the *Drosophila* nerve cord, synaptic density of mechanosensory neurons scales to body size from first to third instar larvae^4^. In the spinal cord of the zebrafish larva, descending neurons lay down tracks chronologically, coinciding with the maturation of swimming behaviours^7^. In the mouse visual circuit, postnatal synaptic remodelling is shaped by intrinsic activity as well as visual stimuli^9^. These and other studies raise the possibility that anatomical changes, from individual synapses to global organization of brain networks^10^, occur. An assortment of genetic and cellular factors have been found to affect morphological and functional maturation of individual synapses^11,12^. Synaptic changes underlie system-level modifications. However, developmental principles for the collective synaptic changes that shape the adult brain are unknown.

Interrogating whole-brain maturation at synapse resolution is difficult. High-resolution electron microscopy (EM) reconstruction is needed to capture structural changes at individual synapses over large volumes^13^. To uncover brain-wide principles of maturation, these methods must be applied to the entire brain, and to brains at different developmental time points. Moreover, multiple animals need to be analyzed to assess structural and behavioural heterogeneity. With recent advances in automation and throughput of EM, this has become uniquely possible using the nematode *C. elegans*, the first animal that allowed the assembly of a complete connectome by serial section EM reconstruction^14,15^.

Serial-section EM has now been used to reconstruct neural circuits with synapse resolution across species^16–22^. But in larger animals, low throughput makes it difficult to acquire whole brain samples and comprehensively assess plasticity. EM has been applied to assess wiring differences between individuals, for example, comparing the pharyngeal circuits of two nematode species^23^, comparing the *C. elegans* male and hermaphrodite connectomes^24^, the effect of genotype or age on the *Drosophila* larval somatosensory^25^ and mechanosensory^4^ circuits, as well as the effect of developmental age on wiring in the mouse cerebellum^26^. These studies examined partial circuits or few samples. The original *C. elegans* connectome was compiled from the EM reconstruction of partially overlapping regions of four adults and an L4 larva. A revisit of this connectome expanded the wiring diagram by re-annotation of original EM micrographs and filled remaining gaps by interpolation^24^, making it more difficult to assess differences between animals.

Here, we leveraged advances in automation and throughput of EM reconstruction to study the brain of *C. elegans* - its circumpharyngeal nerve ring and ventral ganglion - across development. We have fully reconstructed the brain of eight isogenic hermaphroditic individuals at different ages of postembryonic development, from hatching (birth) to adulthood. These reconstructions provide quantitative assessments for the length, shape, and position of every neural and muscle fibre in the nerve ring, as well as of every physical contact and chemical synapse between neurons and muscles, and between neurons and glia. Our quantitative comparisons of these developmental connectomes have revealed organizing principles by which synaptic changes shape the mind of the developing worm.

## Results

### EM reconstruction of eight *C. elegans* brains from birth to adulthood

We developed approaches in ultra-structural preservation, serial ultra-thin sectioning, and semi-automated imaging^27–29^ to reconstruct the connectivity and morphology of all cells in eight individual isogenic hermaphroditic brains of *C. elegans* (N2) at various post-embryonic stages (Fig. S1, 1a, Video 1-2, see Methods). The brain, consisting of the nerve ring and ventral ganglion, includes 162 of the total 220 neurons at birth (L1), and 180 of the total 302 neurons in adulthood of the original connectome^14,30^ (Table S1). The brain also contains 10 glia and synaptic sites of 32 muscles at all stages. We identified every cell across different EM volumes based on their unique neurite morphology and position^14^. Because CANL/R, one pair of cells in the original reconstructions make no synapses in all our datasets, they were excluded from the *C. elegans* connectome. Each neuron was classified as either being sensory, inter, motor, or modulatory (Table S1, Supplementary Information 1, Video 2, see Methods).

In each EM volume, every neuron, glia, and muscle was annotated for chemical synapses to generate a connectome of the brain (Fig. 1b, Fig. S2, Video 2, Supplementary Information 2, see Methods). Chemical synapse annotations include classical synapses, which contain mostly clear vesicles as well as a small number of dense core vesicles, and synapses from modulatory neurons, which contains mostly dense core vesicles (Fig. 1c, see Methods). Presynaptic active zones of chemical synapses were volumetrically reconstructed to determine synapse sizes (Supplementary Information 3). Neuron and muscle processes, but not glia processes, were volumetrically segmented (Supplementary Information 4). Gap junctions were partially annotated (and are shown at http://nemanode.org/), but because they could not be mapped in completion, they were excluded from further analyses.

**Figure 1.**
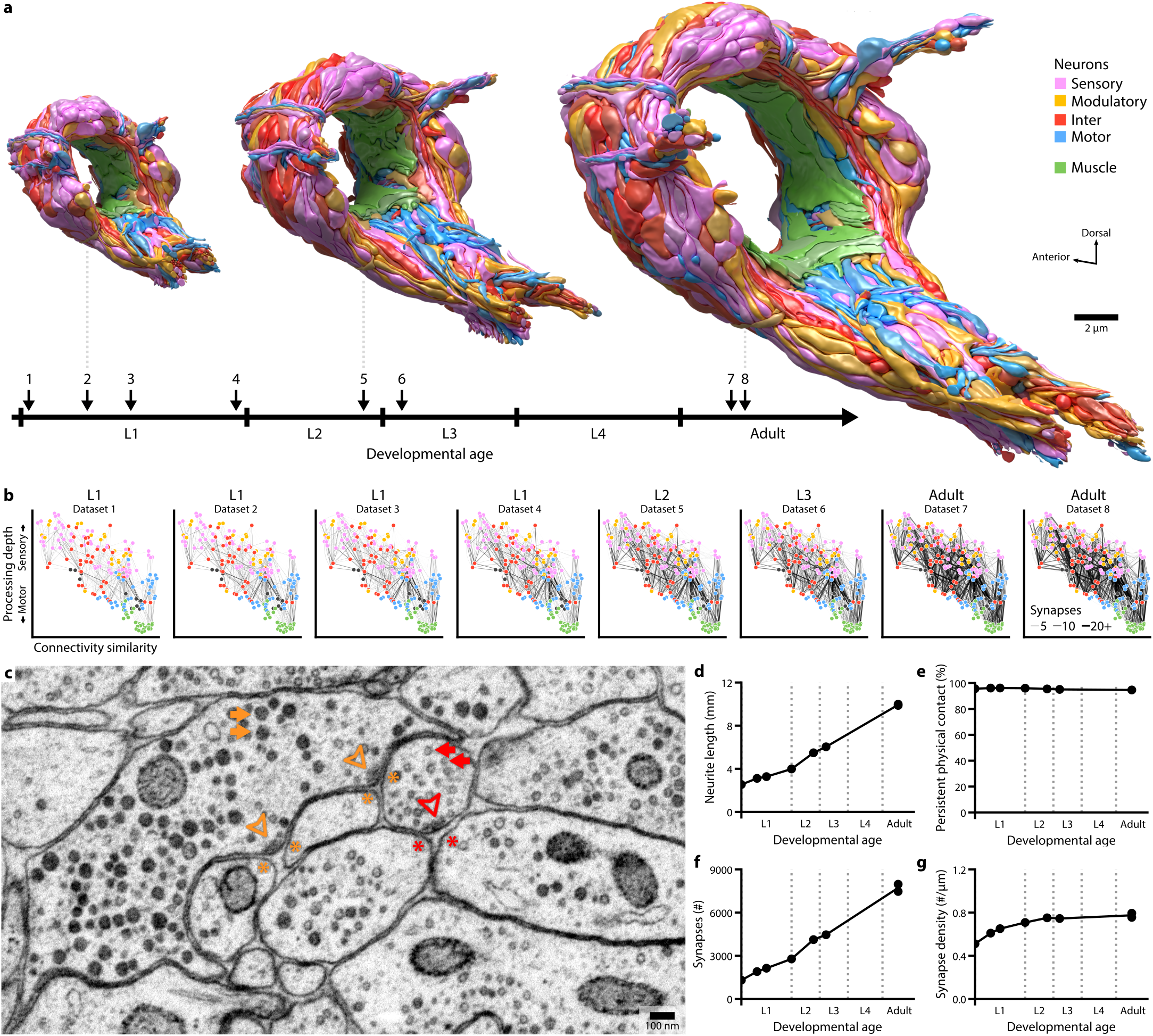
The developing brain maintains its overall geometry. **a**. Developmental timeline of eight reconstructed brains, with volumetric models shown at three stages. Models include all neurites contained in the neuropil, coloured by cell types. **b**. Wiring diagrams for all datasets. Each circle represents a cell. Each line represents a connection with at least one chemical synapse between two cells. The line width indicates synapse number. The vertical axis denotes signal flow from sensory perception (top) to motor actuation (bottom); the horizontal axis denotes connectivity similarity (normalized Laplacian eigenvector 2, see ^15^) where neurons that share partners are positioned nearby each other. Signal flow and connectivity similarity are based on the accumulated connections from all datasets. **c**. A representative EM micrograph of the neuropil (from dataset 3). Presynaptic termini of classical chemical synapses are characterized by a pool of clear synaptic vesicles (red arrows) surrounding an active zone (red arrowhead). Presynaptic termini of chemical synapses of modulatory neurons are characterized by mostly dense core vesicles (orange arrows) distant from the active zone (orange arrowhead). Postsynaptic cells are marked by asterisks. **d**. Summed length of all neurites in the brain exhibits linear increase from birth to adulthood. Each data point represents the total neurite length from one dataset. **e**. Persistent physical contact, the summed physical contact between all neurite pairs at birth that persists across maturation, accounts for nearly all of the contact area at every developmental stage. **f**. Total synapse numbers in the brain exhibits a 6-fold increase from birth to adulthood. **g**. Synapse density, the total number of synapses divided by the summed length of all neurites, is maintained after an initial increase.

We plotted the wiring diagrams conforming to the direction of information flow from sensory perception (Fig. 1b top layer) to motor actuation (Fig. 1b bottom layer). All connectomes are hosted on an interactive web-based platform at http://nemanode.org/. These datasets allow for examination of changes of chemical synaptic connectivity in the context of brain geometry, including the shape and size of each neuron as well as the proximity and contact between each neurite (see below).

### Uniform neurite growth maintains brain geometry

Our volumetric reconstructions revealed striking similarities of brain geometry between developmental stages. The shape and relative position of every neurite in the brain was largely established by birth (Fig. S3a, S3b). From birth to adulthood, the total length of neurites underwent a 5-fold increase (Fig. 1d), in proportion to the 5-fold increase in body length (∼250µm to ∼1150µm). Neurites grew proportionally (Fig. S3b), maintaining physical contact between cells that are present at birth across maturation (Fig. 1e, S3a). Only three neuron classes had changes to their primary branching patterns, each growing a new major branch after birth (Fig. S4, Video 3). Thus, the brain grows uniformly in size without substantially changing the shape or relative position of neurites, maintaining its overall geometry.

In parallel to neurite growth, addition of synapses was extensive even though only a small fraction of physical contacts developed into chemical synapses (Fig. S3c). From birth to adulthood, the total number of chemical synapses increased 6-fold, from ∼1300 at birth to ∼8000 in adults (Fig. 1f). Presynaptic terminals appear as *en passant* boutons, most often apposing the main neurite of a postsynaptic cell. Small spine-like protrusions^14,31^ were postsynaptic at ∼17% of synapses in the adult connectome (Fig. S5a, S5b, Supplementary Information 5). From birth to adulthood, the number of spine-like protrusions increased 5-fold (Fig. S5c), and the proportion of spine-like protrusions apposing presynaptic terminals increased 2-fold (Fig. S5d). Protrusions apposing presynaptic termini were more likely to locate distally along a neurite, whereas protrusions not apposing presynaptic termini were more proximal (Fig. S5e). Spine-like protrusions were found in many neurons and muscles (Fig. S5f, S5g).

Synapse number increased in proportion to neurite length, maintaining a stable synapse density across most developmental stages. The exception was during the L1 stage, when the increase of total synapse number slightly outpaced that of neurite length, leading to increased synapse density (Fig. 1g). This increase coincided with an increasing left-right wiring symmetry (Fig. S6a, S6b). In the adult brain, ∼90% of neurons exist as left-right pairs that mirror one another in position, morphology, as well as connectivity^14^. However, some of these neurons exhibited left-right connectivity asymmetry at birth (Fig. S6a, S6b). The simplest interpretation of this early asymmetry is incompleteness: *C. elegans* hatches before its brain connectivity has been made symmetric, a process which continues by synapse addition during the first larval stage.

### Non-uniform synapse addition reshapes the connectome

From birth to adulthood, addition of synapses both creates new connections and strengthens existing connections. Here, a connection is defined as a pair of cells connected by one or more chemical synapses (Fig. 2a).

**Figure 2.**
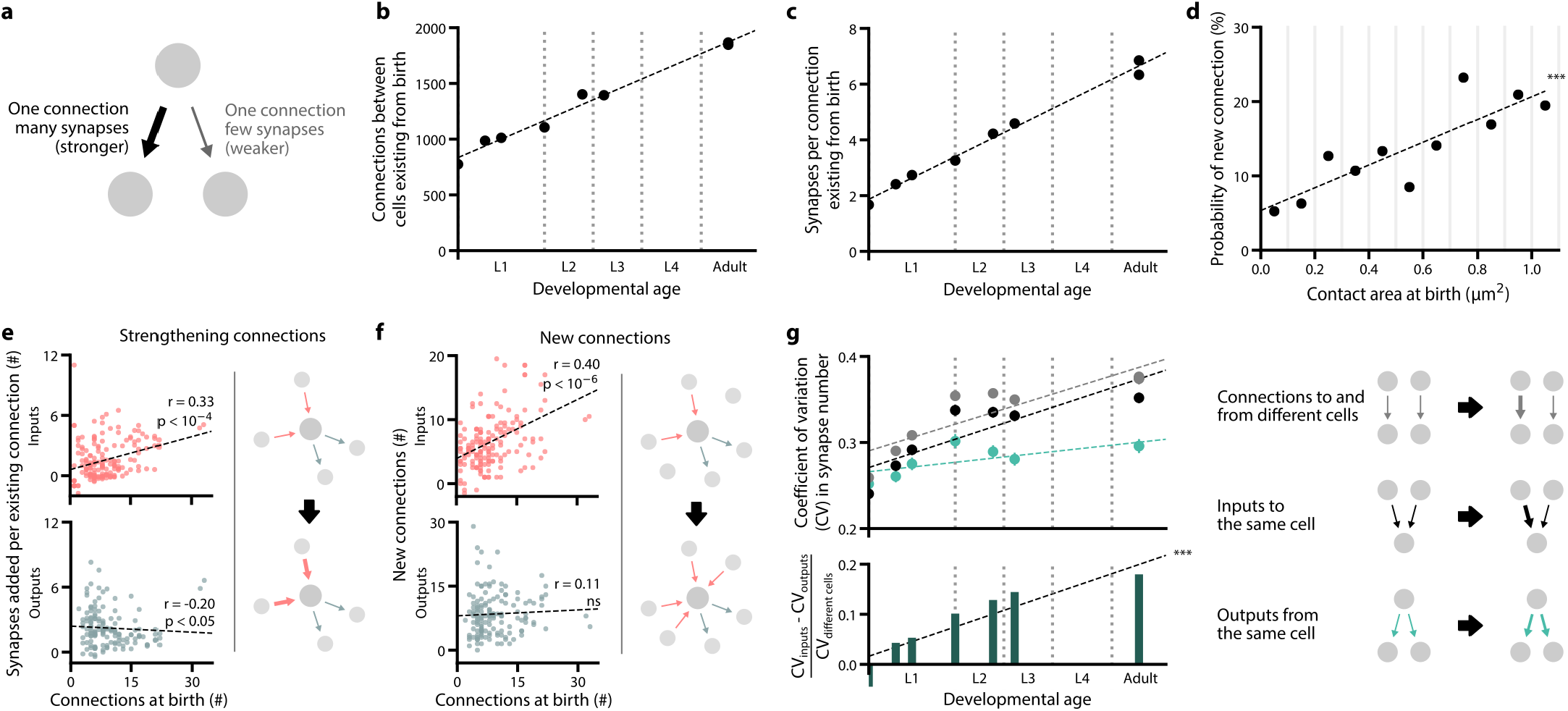
Non-uniform synapse addition reshapes the connectome. **a**. Schematic of two connections. Each connection consists of at least one synapse between two cells. **b**. The total number of connections in the brain between neurons existing from birth exhibits a 2.4-fold increase. **c**. The mean number of synapses per connection existing from birth exhibits a 3.9-fold increase. **d**. The probability of forming a new connection at physical contacts existing from birth. This probability increases with the total contact area between two cells at birth. A new connection is here defined as a connection that appears in adults (datasets 7 and 8) but is absent in early L1 stages (datasets 1 and 2). *** r = 0.87, p = 4.5×10^−4^, Spearman’s rank correlation. **e**. Top: neurons with higher number of connections at birth (dataset 1) are more likely to receive new synapses at existing input connections by adulthood (averaging datasets 7 and 8). Bottom: no positive correlation is observed at existing output connections. Each data point represents one cell. Significance is calculated using Spearman’s rank correlation (top: p = 1.1×10^−5^, n = 166; bottom: p = 0.017, n = 141). **f**. Top: neurons with higher number of connections at birth (dataset 1) are more likely to establish new input connections by adulthood (averaging datasets 7 and 8). Bottom: no correlation is observed at new output connections. Each data point represents one cell. Significance is calculated using Spearman’s rank correlation (top: p = 1.3×10^−7^, n = 166; bottom: p = 0.18, n = 141). **g**. Top: each data point represents the mean coefficient of variation (CV) in the number of synapses for different sets of connections. The CV of output connections from the same cell is maintained. The CV of input connections to the same cell increases over time, at the same rate as connections to and from different cells. Error bars indicate SE. Bottom: the difference between the mean CV for output and input connections relative to connections between different cells grows over time. *** p = 5.3×10^−7^, r = 0.99, Spearman’s rank correlation.

Both synapse and connection number increase during maturation. The 204 cells of the brain were interconnected by ∼1300 total synapses distributed among ∼800 connections at birth (Fig. 2b). Over maturation, addition of synapses strengthened nearly all existing connections. Approximately 4500 synapses were added to connections that were present at birth, such that the mean synapse number per connection increased 4.6-fold, from 1.7 synapses per connection at birth to 6.9 by adulthood (Fig. 2c). In addition, many new connections formed. Approximately 1200 synapses formed between previously nonconnected neurons resulting in a 2.4-fold increase in total number of connections between cells present at birth (Fig. 2b).

Synapse addition did not occur uniformly across the brain from birth to adulthood. We found preferential synapse addition in multiple contexts.

First, new connections were more likely to form at physical contacts between neurons with large contact areas at birth (Fig. 2d). Physical contacts formed at birth therefore appear to create a constant scaffold within which network formation unfolds.

Second, synapse addition preferentially strengthens inputs to neurons with more connections at birth. At birth, some neurons already had far more connections than others (Fig. S6c). These neurons, which we refer to as hubs, disproportionately strengthened their existing input connections over time (Fig. 2e). Hubs also disproportionately established more new input connections over time (Fig. 2f). Interestingly, hub neurons did not disproportionately increase their outputs (Fig. 2e, 2f). The increase in inputs was evident even for neurons with only more output connections at birth (Fig. S6d, S6e). Thus, during maturation the flow of information is progressively focused onto the most highly-connected neurons at birth.

Third, synapse addition selectively strengthened a cell’s individual connections. We found that there was no correlation in the strengthening of existing input connections to each cell from different presynaptic partners (Fig. S6f), leading to a divergence in the relative strengths of different inputs (Fig. 2g). However, strengthening of the existing output connections from each cell were correlated (Fig. S6f), maintaining their relative strengths (Fig. 2g). Thus, it appears that each cell regulates the strengthening of its own outputs but does not dictate the relative strengthening of its inputs.

Lastly, in contrast to mammals where pruning is a hallmark of early nervous system development, we did not observe systematic synapse elimination in *C. elegans*. Synaptic connections are not often removed; remodeling that diminishes synaptic connections is mediated by selective strengthening of other connections.

### Isogenic individuals have both stereotyped and variable connections

We mapped the change in synapse number for each connection across developmental stages to evaluate the change in connection strength. Using these maps, we classified each connection as either stable, developmentally dynamic, or variable (Fig. 3a, S7, Supplementary Information 6, see Methods).

**Figure 3.**
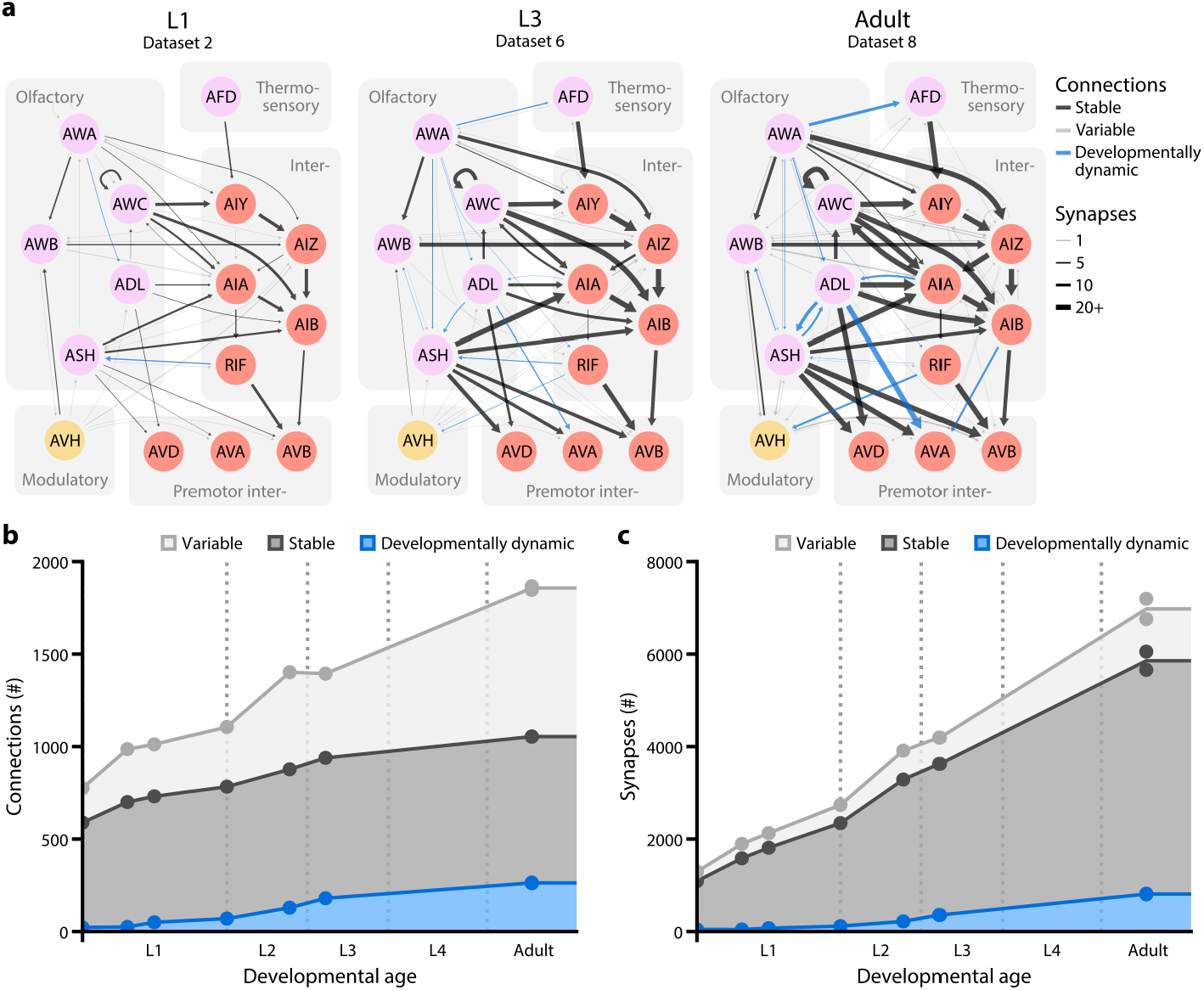
Isogenic individuals have both stereotyped and variable connections. **a**. A sensory circuit across maturation. Left: L1 (dataset 2), center: L3 (dataset 6), right: adult (dataset 8). Circles represent cells, colour-coded by cell types. Colour-coded lines represent stable (black), developmentally dynamic (blue), and variable (grey) connections. Line width represents synapse number. **b**. The total number of stable, developmentally dynamic, and variable connections in each dataset. **c**. The total number of synapses that constitute stable, developmentally dynamic and variable connections in each dataset.

Stable connections were present from birth to adulthood and maintained their relative strength in proportion to one another. Developmentally dynamic connections significantly increased or decreased their relative strength in a stereotyped manner, sometimes forming new connections or more rarely eliminating existing connections at specific life stages. Variable connections exhibited no consistent trend in their changing strength, and were not present in every animal.

In the adult connectome, stable and variable connections each represented ∼43% of the total number of connections, whereas developmentally dynamic connections represented ∼14% (Fig. 3b). We observed similar partitions when connections were classified by changes in either synapse size (Fig. S8a) or synapse density (Fig. S8b), instead of synapse number, suggesting that synapse number (see Methods - Connectome annotation) can be a good proxy for synapse size.

Stable connections contained more synapses than variable ones (6.6±5.8 versus 1.4±1.0 synapses per connection, respectively, in adult), and thus constituted a larger proportion (∼72%) of total synapses (Fig. 3c). Nonetheless variable connections were surprisingly common. Like other connections, variable connections formed at existing and maintained cell contacts with little exception (Fig. S8c, also see Fig. S3a, S7). The number of variable connections in the adult (∼800) is similar to the number of stable connections (∼800). The total number of synapses that constitute variable connections in the adult (∼1100) is even greater than that of developmentally dynamic connections (∼800). Synapses that comprise variable connections were comparable in size to those of stable connections, and were similarly distributed between monoadic and polyadic synapses (Fig. S8d-S8g).

Moreover, not all variable connections consisted of few synapses and not all stable connections consisted of many synapses (Fig. S8h). The range of synaptic strength of stable and variable connections makes it difficult to set them apart by thresholding. Any threshold to filter postsynaptic partners – by synapse number, synapse size, or number of postsynaptic partners – excluded both variable and stable connections (Fig. S8f-S8i).

We considered the possibility that variability is more prominent during development than in the mature connectome. A conservative measure of variability in the adult stage can be made by comparing our two adult datasets and the original adult connectome^14^. When using these adult datasets to quantify variability, variable connections still made up ∼50% of all connections (Fig. S9a, S9b). Thus, variable connections are prominent in the *C. elegans* connectome.

### Variable connections are not uniformly distributed among cell types

To visualize the distribution of different classes of connections, we separately plotted their occurrences in the wiring diagram (Fig. 4a). Stable and developmentally dynamic connections represent the portion of the connectome that is shared across animals. Variable connections represent the portion that is unique to each animal.

**Figure 4.**
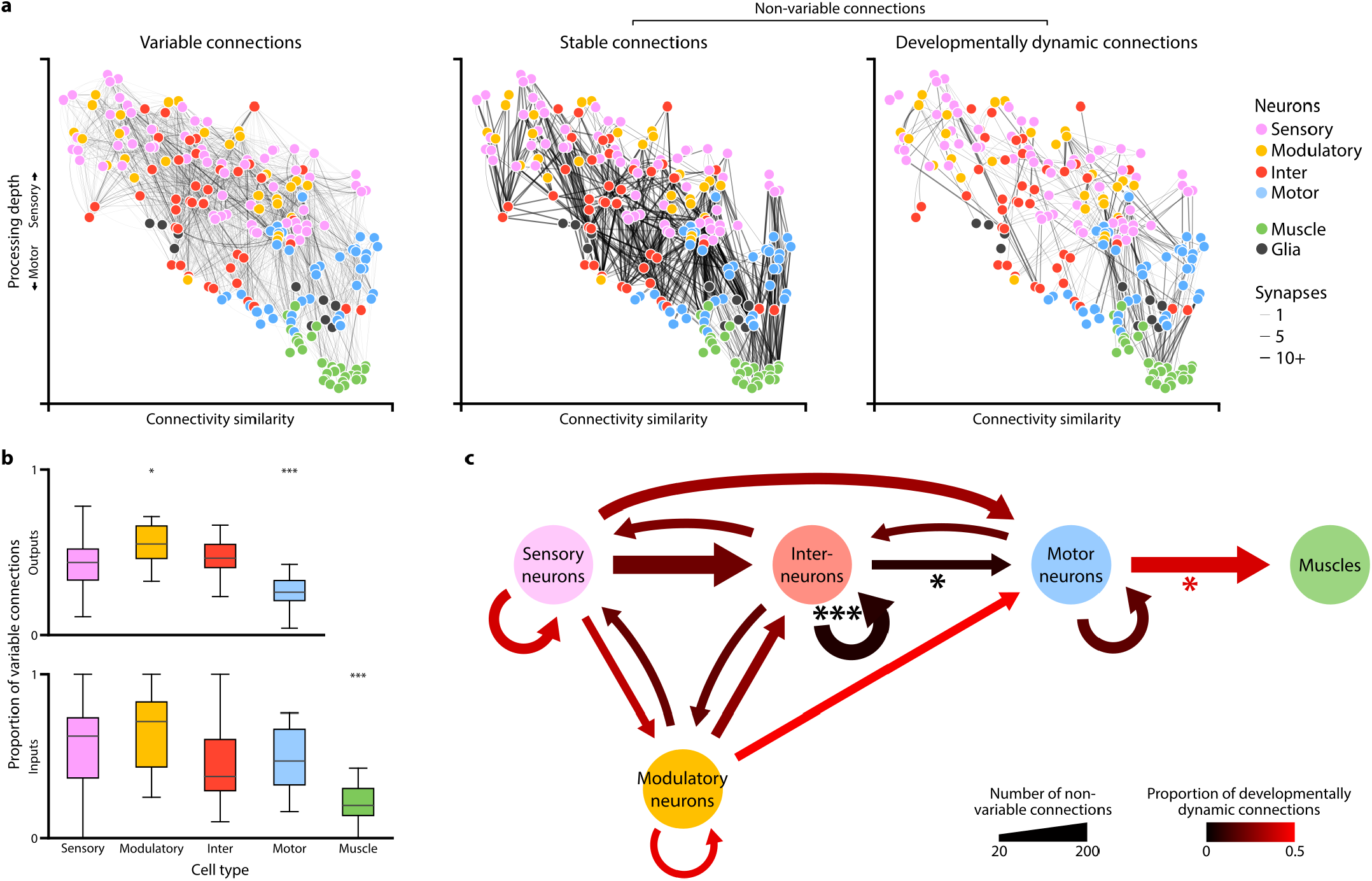
Non-uniform distribution of variable and developmentally dynamic connections. **a**. Wiring diagrams for variable, stable, and developmentally dynamic connections. Each line represents a connection observed in at least one dataset. Line width indicates the largest number of synapses observed for a connection across datasets. Each circle represents a cell. Cell coordinates are represented as in Fig. 1b. **b**. Comparison of the proportion of variable and non-variable connections for each cell type. Non-variable connections include stable and developmentally changing connections. Cell types with significantly higher or lower proportions of variable connections are indicated, ** p < 10^−2^, *** p < 10^−3^, n = 22-57, Mann–Whitney U test, FDR adjusted using Benjamini–Hochberg correction. Center line, median; box limits, upper and lower quartiles; whiskers, 1.5x interquartile range; outliers not shown. **c**. Wiring diagram showing non-variable connections between different cell types. Line width indicates the number of connections. Line color indicates the proportion of developmentally dynamic connections. Lines with significantly different proportions of developmentally dynamic connections are indicated, * p < 4.1×10^−2^, *** p = 2.0×10^−5^, two-tailed Z-test, FDR adjusted using Benjamini–Hochberg correction (n_inter-inter_ = 160, n_inter-motor_ = 52, n_motor-muscle_ = 145).

We quantified the proportion of variable connections in the inputs and outputs of each cell type (Fig. 4b). Modulatory neurons had significantly higher amounts of variability in their output connections than other cell types, whereas motor neurons had significantly less (Fig. 4b upper panel). Consistent with the lowest variability in motor neuron output, muscles exhibited the lowest variability in their inputs (Fig. 4b lower panel).

The higher prevalence of variable connections between certain cell types remained evident when variable connections were defined only among adult datasets (Fig. S9c) and when connections with fewer synapses were excluded (Fig. S8i). The low variability of connections from motor neurons to muscles could not be simply explained by saturation of their physical contacts by synapses (Fig. S9d). We also considered that neurons with more synapses may exhibit more stochastic synapses or have more annotation errors. However, the proportion of variable connections did not scale with the number of synapses (Fig. S9e, S9f).

Rather, the likelihood of a neuron to generate variable connectivity is likely a property of its cell type. The high stereotypy of synapses from motor neurons to muscles may reflect a requirement for high fidelity in circuits for motor execution. Modulatory neurons, which may secrete monoamines and neuropeptides by volume-release, may have the weakest requirement for precise spatial positions of synaptic outputs because they exert long-range effects.

### Interneuron connections are stable during maturation

Excluding variable connections allows us to assess shared developmental connectivity changes across animals. We found that developmentally dynamic connections were not uniformly distributed among cell types or circuit layers (Fig. 4c). Connections between interneurons and from interneurons to motor neurons had disproportionately more stable connections than developmentally dynamic connections (Fig. 4c). In contrast, all other connections between and from sensory, modulatory, or motor neurons had many developmentally dynamic connections. This trend remains evident when developmentally dynamic connections were classified by synapse size instead of by synapse number (Fig. S10a middle panel). Developmentally dynamic connections were particularly prevalent from motor neurons to muscles. Each motor neuron progressively increases its connections with more muscles in a stereotypic pattern (Fig. S7). The abundant but high stereotypy of this developmental connectivity change means that motor neurons exhibit the lowest proportion of variable connections (Fig. 4b upper panel). Developmentally dynamic connections were also prevalent between many sensory neurons, and from sensory neurons to interneurons and motor neurons (Fig. 4c, Fig. S7). Spine-like protrusions may facilitate developmental connectivity changes, as developmentally dynamic connections were twice as likely to involve spine-like protrusions than stable and variable connections (**??**).

These findings show that maturation changes how sensory information is integrated and relayed to downstream neurons. Maturation also changes motor execution. However, the layout of interneuron circuits, the core decision-making architecture of the brain, is largely stable from birth to adulthood.

### Increase in both feedforward signal flow and modularity across maturation

With connectomes of complete brains, we were able to ask how the sum of synaptic changes leads to collective changes in information processing across maturation.

First, we examined how synaptic changes affect information flow in the brain. The directionality of signal flow between cells can be viewed as either feedforward, feedback, or recurrent (Fig. 5a). We classified connections that constitute synapses from the sensory to motor layer as feedforward, connections from the motor to sensory layer as feedback, and connections between neurons of the same type as recurrent. Among stable connections, synapse addition strengthened existing feed-forward connections more than feedback or recurrent connections (Fig. 5b). We observed the same trend when considering synapse size instead of synapse number (Fig. S10b). This difference was not simply due to increased inputs onto stable interneuron circuitry, as interneuron connections exhibited a similar increase in synapse number compared to connections for sensory inputs and motor outputs (Fig. S10a right panel). The addition of developmentally dynamic connections also preferentially increased feedforward signal flow (Fig. 5c). In contrast, developmentally dynamic connections that were weakened across maturation tended to be feedback and recurrent. Cumulatively, the proportion of synapses that constitute feed-forward connections gradually increased (Fig. 5d). Thus, one global pattern of brain maturation augments signal flow from sensation to action, making the brain more reflexive (and less reflective) with age.

**Figure 5.**
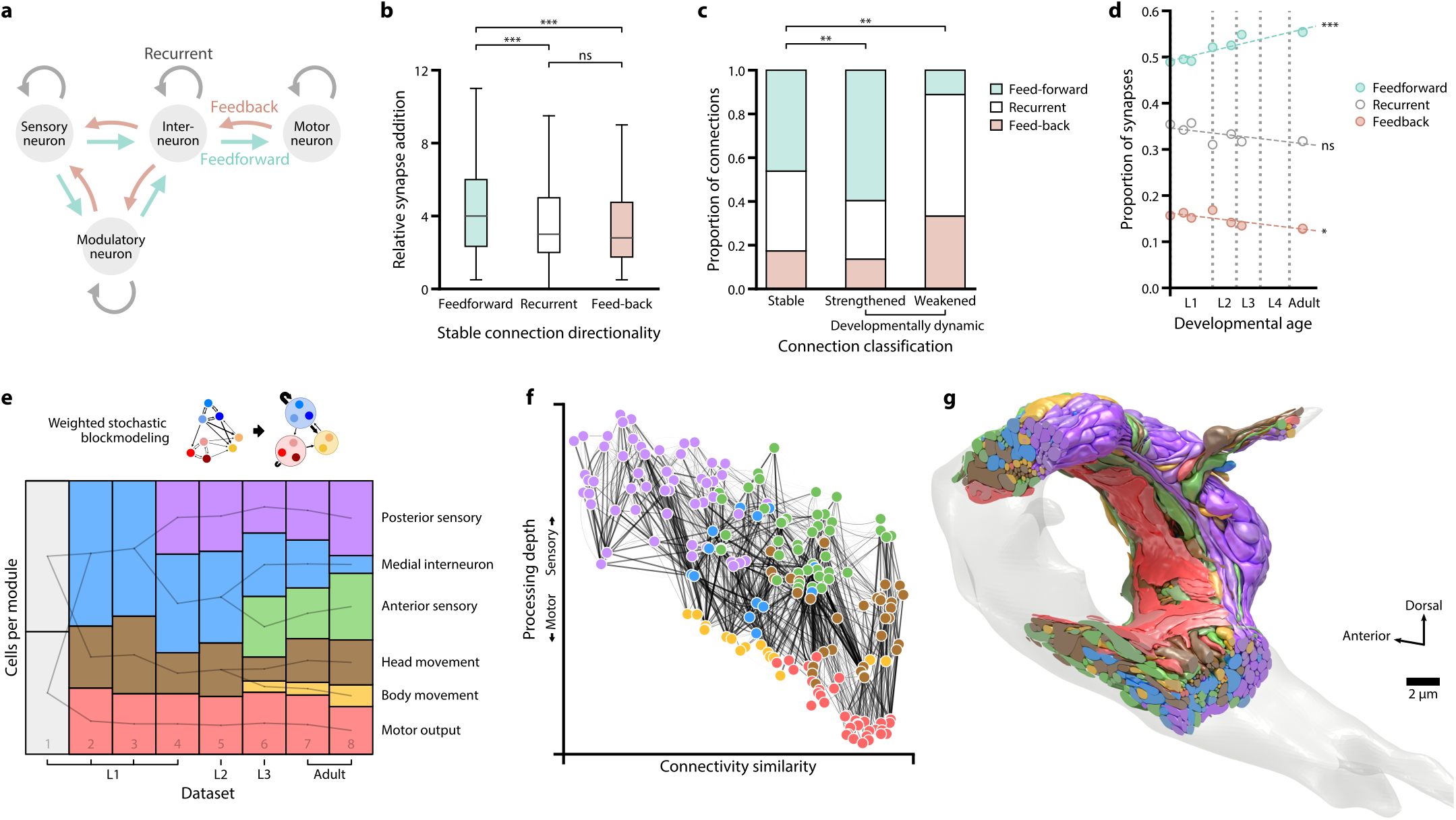
Increase in both feedforward signal flow and modularity across maturation. **a**. Schematic of feedforward, feedback, and recurrent connections defined by cell types. **b**. The number of synapses for stable connections in adults (datasets 7 and 8) relative to birth (datasets 1 and 2). Stable feedforward connections are strengthened more than stable feedback and recurrent connections. ns (not significant) p = 0.13, *** p < 0.001, Mann–Whitney U test, FDR adjusted using Benjamini–Hochberg correction (n_feedforward_ = 301, n_recurrent_ = 229, n_feedback_ = 107). Center line, median; box limits, upper and lower quartiles; whiskers, 1.5x interquartile range; outliers not shown. **c**. Proportions of feedforward, feedback, and recurrent connections for stable and developmentally dynamic connections. ** p < 0.005, two-tailed Z-test of the proportion of feedforward connections, FDR adjusted using Benjamini–Hochberg correction (n_stable_ = 737, n_added_ = 198, n_weakened_ = 18). **d**. Proportions of the total number of synapses in feedforward, feedback, and recurrent connections. ns (not significant) p = 0.11, * p = 0.017, *** p = 2.0×10^−4^, Spearman’s rank correlation, FDR adjusted using Benjamini–Hochberg correction. **e**. The number of cells in each module across maturation, determined by weighted stochastic blockmodeling. Modules connected by a line share significant number of neurons. See Table S2 for cell membership of each module. The motor output module comprises head muscle cells that are part of the brain connectome; the head movement module comprises motor neurons that innervate and coordinate head muscle cells; the body movement module comprises descending and premotor interneurons that regulate activities of body wall muscles; the anterior sensory module comprises labial sensory neurons, the posterior sensory module comprises amphid sensory neurons; the medial interneuron module comprises the remaining sensory neurons and the majority of interneurons (see Table S2). **f**. The wiring diagram for the adult connectome (dataset 8), with each cell colored by its assigned module. Cell coordinates are represented as in Fig. 1b. **g**. A 3D model of the adult brain (dataset 8), with each cell colored by its assigned module.

Next, we examined the community structure of the brain. We used weighted stochastic blockmodeling (WSBM) to group neurons of similar connectivity into distinct modules^32^. At the adult stage, the modularity corresponds to six congregations of cells with distinct functions (Fig. 5e, 5f, Fig. S11, Table S2). Sensory and interneurons separate into three modules: anterior sensory (including labial sensory neurons), posterior sensory (including amphid sensory neurons and associated interneurons), and medial interneuron (including other sensory neurons and the majority of interneurons). Head motor neurons and descending premotor interneurons for body movement separate into two modules. Muscle cells constitute another module.

When we measured modularity at earlier developmental stages, we discerned progressively fewer modules (Fig. 5e, Fig. S11, Table S2). At birth, WSBM identified two discernible modules. Most of the increase in discernible modularity is due to a smaller fraction of total synapses over development. 74% of new synapses are added to stable connections without increasing modularity (Table S3, bottom row). The difference in discernible modularity is mostly attributed to developmentally dynamic connections, which only represent 14% of new synapses (Table S3, middle row). Variable connections, which are not uniformly distributed among cell types, also contributed to module segregation (Table S3, top row). The increased connectivity increases the number of discernible modules of closely connected neurons in the adult brain (Fig. 5g, Video 4). The physical proximity of neurons in these modules is reminiscent of distinct brain lobes with different behavioral roles.

## Discussion

To learn the emergent principles from studying synaptic changes of an entire brain across maturation, we analysed eight isogenic *C. elegans* beginning with the earliest larva stage and ending with the adult. Previous lineage studies revealed that the vast majority of post-embryonic neurogenesis and differentiation occurs during the L1 and L2 stages^30^. We reconstructed three L1s, two L2 and one L3 animals at six different developmental time points, to afford the temporal resolution in capturing continuous connectomic changes during the period of most rapid growth. We reconstructed two adults to make direct comparisons between animals of the same age and with the original published connectome. While it took more than a decade to assemble the first *C. elegans* connectome^14^, the advent of automation in sample sectioning, image acquisition, and data processing sped up the process, allowing our complete brain reconstructions of multiple animals in less time.

We found that several general features remained largely unchanged from the earliest larva to the adult stage. For example, the overall geometry of the brain, the three-dimensional shape, relative placement of individual neurons, and their physical contacts was surprisingly stable. Established neurite neighbourhoods^33^ at birth provides the structural platform that both constrain and support wiring maturation.

In contrast, the total volume of the brain enlarged about 6-fold. However, changes in brain connectivity were not simply explained by enlargement of existing structures. While there was a 5-fold increase in the number of synapses, these synaptic changes were not distributed uniformly through the network. Rather, they appeared to be organized by several developmental principles that profoundly shape how the brain’s network changes.

The principles that we uncovered are illustrated in Fig. 6 and listed below. At one level, we observed patterns of synaptic remodeling that differentially alter the number and strength of connections, applied to every neuron. At a second level, we observed patterns of synaptic remodeling that differ between cell types (i.e., sensory neurons, modulatory neurons, interneurons, and motor neurons). At the third level, we observed network-level changes that alter the directionality of information flow and the segregation of information processing throughout the brain. We propose that these three levels of synaptic remodeling (listed below) might contribute to the behavioral maturation of the growing animal.

**Figure 6.**
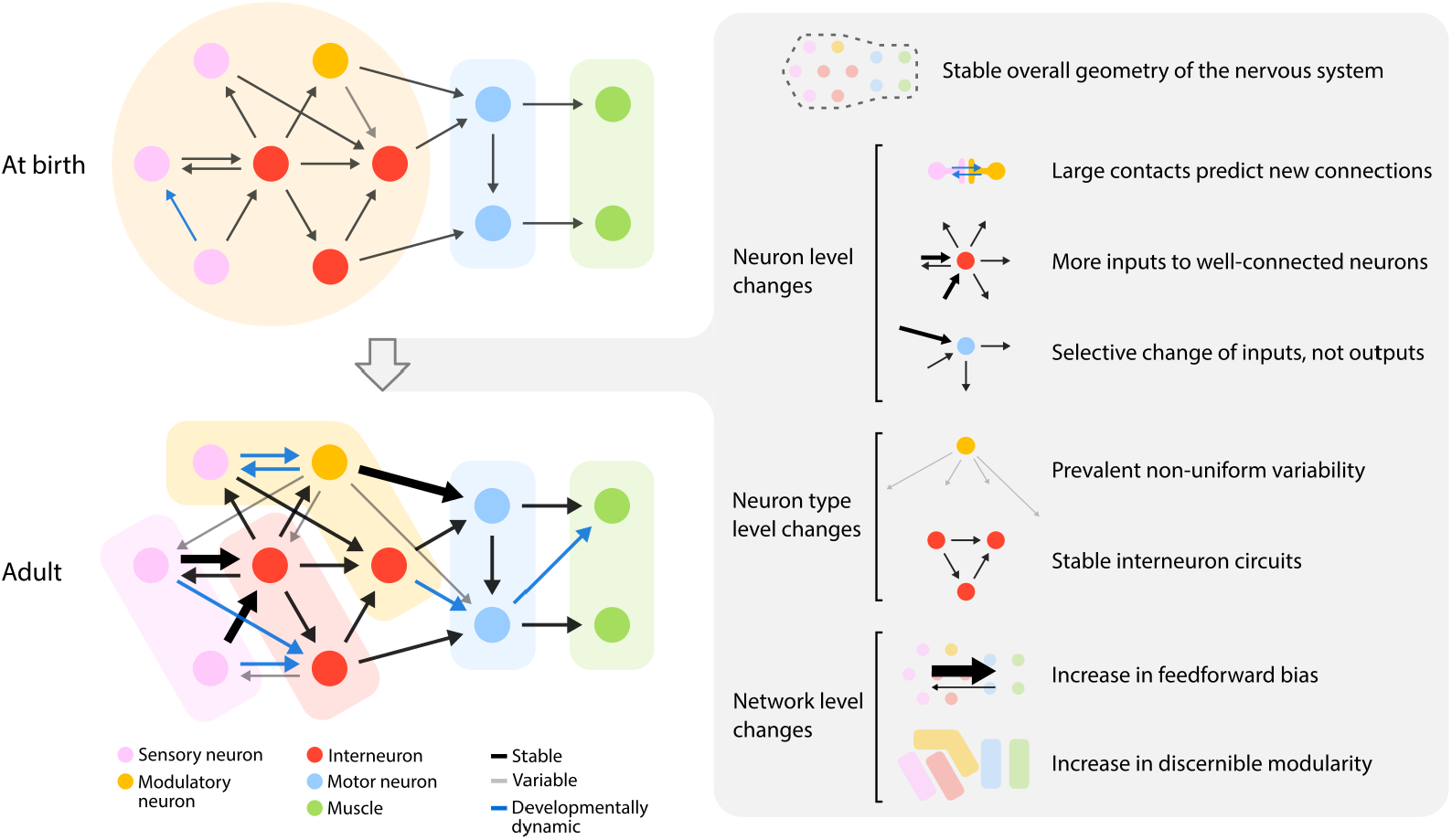
Principles of connectivity changes across maturation. Left: schematic of brain-wide synaptic changes from birth to adulthood. Right: emerging principles of maturation that describe synaptic changes at the level of brain geometry, individual neurons, neuron types, and entire networks. Thicker lines represent stronger connections with more synapses.

### Large contacts predict new connections

Because the overall geometry of the brain is constant, physical contacts between neurites from birth to adulthood are invariant with little exception. Nearly all new synapses appear at sites where these physical contacts already exist, both adding synapses to connections between neurons and creating new connections between neurons. The larger the physical contact, the greater the probability of a new connection. Therefore, the brain’s geometry at birth creates the scaffold upon which adult connectivity is built.

### More inputs to well-connected neurons

We found that developmental synapse addition was not equal among all neurons. Cells with larger numbers of connections at early stages receive disproportionately more new synapses, both strengthening existing input connections and creating new input connections. In contrast, these neurons see less synapse addition to their output connections. Thus, well-connected neurons become better integrators of information, but not broader communicators of that information.

### Selective change of a neuron’s inputs, but not outputs

We also found a pattern in how synapses selectively change the strengths of existing connections. The strength (synapse number) of input connections that converge on the same neuron tends to become more heterogeneous. In contrast, the outputs from the same neuron maintain their relative strengths. Neurons thus become differentially driven by a subset of their presynaptic partners, but distribute that information uniformly among their postsynaptic partners.

### Prevalent variability in the connectomes between animals

Each animal has connections between neurons that are not found in other individuals. These variable connections tend to consist of small numbers of synapses, but represent almost half of connected neuron pairs in the mature brain. This variability is most prominent among modulatory neurons and least prominent among motor neurons.

### Stable interneuron circuits

We discovered remarkable stability in the wiring between interneurons that may constitute the core decision-making architecture of the developing brain. In contrast, there are extensive developmental wiring changes among other cell types.

### Increase in feedforward bias

At the level of the entire network, we discovered a change in the directionality of information flow. Synaptogenesis over development preferentially creates new connections and strengthens existing connections in the direction of sensory layers to motor layers. This makes the network more feedforward and reflexive over time.

### Increase in discernible modularity

Synaptogenesis increases the discernible modular structure of the brain, making it possible to increasingly resolve sub-networks for sensory or motor processing with maturation.

These principles have ontogenetic, phylogenetic, and functional implications discussed below.

#### The *C. elegans* wiring diagram is not stereotyped

We found considerable variability in chemical synaptic connectivity in our set of isogenic animals. In contrast, the extent of physical contacts between neurites at birth were maintained across developmental stages (Fig. 1g and Table SX). About 43% of all cell-cell connections, which account for 16% of total number of chemical synapses, are not conserved between animals. This degree of variability contrasts with the view that the *C. elegans* connectome is hardwired. The idea that individual neurons have identical connectivities probably stemmed from the finding that the same *C. elegans* neuron is identifiable in each animal by virtue of its mostly stereotyped lineage and morphology^14,30,34^. This stereotypy implies that many properties are genetically determined. The reasoning was that if genetic regulation is strong, an isogenic population is more likely to exhibit stereotyped connections between cells. The original connectome, which consisted of compiled annotations from two complete nerve rings - JSH (L4 larva) and N2U (adult), and one partial nerve ring N2T (adult) - did not address variability^14^.

We found that variable connections on average contained fewer synapses than non-variable connections. Interestingly, partial reconstructions of the mammalian thalamus suggests that weaker connections may correspond to incidental wiring^19^.

We also found that synaptic variability between animals is not uniform among cell types. For example, modulatory neurons have considerable variability in their output connections whereas motor neurons have little variability in their outputs. This contrast suggests that variability is in some way regulated between cell types, and may therefore be genetically determined and functionally important. For example, behavioural variability between animals can confer a fitness advantage to a population^35^. Synaptic variability may be a source of such behavioural variability, e.g., in the *Drosophila* visual system, variability among neurite morphologies has been linked to behavioural individuality^36^.

Despite being isogenic, *C. elegans* exhibits individual variability in its behaviors^37^ which could be related to the synaptic variability we describe. One mechanism that might give rise to synaptic variability may be differences in gene expression^38^. Stochastic variability of expression levels has been observed even in the housekeeping genes in *C. elegans* embryos^39^. Neuronal activity can also be a driving force for synaptic remodeling. Individuals from an isogenic population reared in similar conditions will still experience differences in their local environments throughout life, a source of differences in neuronal activity that may translate into wiring variability in the fruitfly^40^. In *C. elegans*, L1 and adult animals have been shown to have differences in their olfactory behaviors^41^. Adult olfactory behaviors can also be modified by the early olfactory experiences of the L1 larva^42^.

Variability in the placement of individual synapses between neurons in the context of largely stereotyped nervous systems has been observed in other small invertebrates. EM reconstruction of isogenic *Daphnia maga* revealed both stereotyped and variable synapses in their optic lobes^43^. EM reconstruction of the visual systems of two closely related *Platyneris* larvae also revealed both stereotyped and variable synapses^20^. The original connectome of *C. elegans* was also examined for inter-animal variability by comparing the nerve ring connectomes of the JSH series (an L4 animal) and N2U^44^. They noted that the numbers of synapses between connected neurons were more variable between animals than between the left and right sides of the same animal. Consistent with our observations, they noted that connections between neurons in JSH and N2U with fewer than three synapses could also be variable between these original datasets. With our eight datasets, we have been able to quantify the patterns of variable and stereotyped synapses and synaptic connections across cell types and across development. We note that the intrinsic variability in the number of synapses between neuron pairs may partly explain previous observations using in vivo fluorescence-labeling labeling of pre- and postsynaptic markers. In a study of synapse formation in the motor circuit, for example, the numbers of fluorescent puncta corresponding to pre- and postsynaptic markers differed across life stage and between animals^45^. Some of this variability in light microscopy of synaptic puncta may be due to animal-to-animal variability in synapse formation that we and others have observed using serial section EM.

#### Developmental changes in the periphery of the connectome versus constancy in the central core

Why is interneuron connectivity more stable across maturation when compared to the sensory input and motor output of the brain? From an evolutionary standpoint, it may not be surprising that the part of a nervous system that physically interacts with the outside world, the sensory and motor systems, is under high evolutionary pressure to maintain an animal’s fitness in changing environments. Such evolutionary changes in the nematode brain (phylogeny) may have accrued as developmental changes (ontogeny) in its wiring diagram.

Stability of the core parts of the nervous system across maturation implies that the central processing unit is robust enough to be used in different contexts. Maturation changes the flow of sensory information into the central processor and the readout of motor execution from the central processor, without changing the central processor itself. Sensory maturation may reflect changes caused by learning and memory^42^. Motor circuit maturation may reflect adaptations to the changing musculature of the growing animal body^46^.

#### The connectome becomes more feedforward during maturation

We observed an increased feedforward-bias of the adult brain that may be more effective in rapid information processing and making reflexive decisions. In contrast, the juvenile network with more feedback connections may have a greater capacity for learning and adaptation. Interestingly, feedback is also what is used to train some artificial neural networks that perform machine learning. After these artificial networks achieve desired performance, they operate in a feedforward manner^47^. The architecture of the adult nematode brain may be a consequence of feedback-mediated optimization of its sensorimotor pathways.

#### The modularity of the adult connectome

We observed an increase in the discernible community structure of the brain’s network with age. With increased numbers of synapses and connections in the adult brain, it becomes possible to resolve functional communities among neurons that are physically close to one another (Fig. 5g). These communities form spatially compact areas for sensory or motor processing, reminiscent of distinct brain areas in larger animals.

#### Perspectives

In larger animals that mature more slowly, maturation involves extensive changes in the nervous system. Apoptosis, neurite degeneration, and synapse pruning remove unwanted circuitry^48^. Neurogenesis, neurite growth, and synapse formation create new circuitry^49^. For the short-lived *C. elegans*, maturation must be fast and efficient. In its small nervous system, each cell is unique, thus each may be characterized by an intrinsic propensity for synaptic remodeling. These changes occur in the context of its stable morphology and fixed amount of physical contact between neighbouring neurites. With these constraints, the nematode has evolved a broad set of principles for synaptic maturation to build its adult brain (Fig. 6).

Connectome comparisons have revealed instances of wiring plasticity caused by development or genetics. In the *Drosophila* larva, the mechanosensory circuit at two different larval stages is maintained by scaled synapse growth^4^. In the mouse, activity-driven connectivity changes have been uncovered in the cerebellum^26^. Differences in the pharyngeal circuits of different nematode species point to genetic specification of wiring patterns^23^. Comparison of the *C. elegans* male and hermaphrodite reveals sexual dimorphism in their nervous systems with different numbers of neurons and shared and divergent connections^24^.

Future work will extend the study of the development of the *C. elegans* connectome. First, we have not included gap junctions, critical components of the nervous system, in our analysis. Our online connectome database (www.nemanode.org) includes electrical synapses where gap junctions were most visible. But improvements in sample preparation and analysis are needed to reach the same level of confidence and throughput as we reached for chemical synaptic networks throughout development. Second, we have analyzed only one connectome at most time points. This allowed us to compare stable, variable, and developmentally dynamic synaptic networks across development but not to assess animal-to-animal variability at each age. Increased throughput and the analysis of many animals at each age will allow analysis of the statistical properties of synaptic connectivity.

In the *C. elegans* brain, synaptic remodeling leads to changes from the cell to network level, with likely functional consequences on behaviour. Most investigations of flexibility in neural circuits and behaviours focus on functional modulations of connectomes that are assumed to be anatomically static^50,51^. Our comparison of connectomes argues that the maturation and variability of brain and behaviour are not separated from wiring changes. Moreover, comparative connectomics is needed to understand the origin of similarities and differences in structure and behaviour, within and across species. High-throughput and high-resolution electron microscopy are necessary to establish the foundation for understanding how genes, experience, and evolution create the behaving adult.

## Methods

### Data acquisition

We studied wild-type (Bristol N2) animals reared in standard conditions: 35×10mm NGM-plates, fed by OP50 bacteria, and raised at 22.5 °C^52^. The animals were within a few generations of the original stock acquired from *Caenorhabditis elegans* Genetics Center (CGC) in 2001. All samples used in this study were derived from three batches of EM preparation.

Each EM sample was prepared and processed as previously described^29^ with small modifications to the substitution protocol of the last 3 datasets (protocol in preparation). In short, isogenic samples reared in the same environment were highpressure frozen (Leica HPM100 for datasets 1-5 and Leica ICE for datasets 6-8) at different stages of post-embryonic development. High-pressure freezing was followed by freeze-substitution in acetone containing 0.5% glutaraldehyde and 0.1% tannic acid, followed by 2% osmium tetroxide. For each life stage, we selected animals based on their overall size and morphology for EM analysis. The precise developmental age of each larval animal was determined based on its cellular compositions relative to its stereotyped cell lineage^30^, as well as the extent of its neurite growth (see Supplementary Information 7). Three samples (datasets 2, 6, and 7) were prepared for transmission electron microscopy (TEM). Five samples (datasets 1, 3, 4, 5, and 8) were prepared for scanning electron microscopy (SEM).

For TEM, samples were manually sectioned at ∼50nm using a Leica UC7 ultramicrotome, collected on formvar-coated slot grids (Electron Microscopy Sciences, FF205-Cu), post-stained with 2% aqueous uranyl acetate and 0.1% Reynold’s lead citrate, and coated with a thin layer of carbon. Images were acquired using an FEI Tecnai 20 TEM and a Gatan Orius SC100 CCD camera.

For SEM, samples were serial sectioned at ∼30-40nm and collected using an automated tape-collecting ultramicrotome (ATUM)^53^. The tape was glued to silicon wafers, carbon coated, and sections post-stained with 0.5% uranyl acetate (Leica Ultrostain I, Leica Microsystems) and 3% lead citrate (Leica Ultrostain II, Leica Microsystems). Images were collected semiautomatically using custom software guiding an FEI Magellan XHR 400L^54^.

All images were acquired at 0.64-2 nm/px (∼25,000x). In total, these datasets comprise 94374 images, 5 teravoxels, and 2.4×10^5^ µm^3^. Images were aligned using TrakEM2^55,56^ and imported into CATMAID^57^ for annotation.

### Connectome annotation

All cells within the brain were manually reconstructed by skeleton tracing in CATMAID^57^. The brain was defined as the nerve ring and ventral ganglion neuropil anterior of the ventral sublateral commissures. Chemical synapses and gap junctions were mapped manually. Chemical synapses were mapped fully and gap junctions partially. To reduce biases from different annotators, for chemical synapses, all datasets were annotated independently by three different people; only synapses that were agreed upon by at least two independent annotators were included in the final dataset.

The same neurons were unambiguously identified in all datasets based on their soma position, neurite trajectory, and stereotypic morphological traits, as described^14^. In the original connectome datasets, as well as ours, some variability in cell body position and neurite trajectory was observed (see Supplementary Information 8). However, every cell could still be unambiguously identified in every dataset because the combined anatomical features and neighbourhood for each cell is unique. Negligible amounts of neuropil in our reconstructions could not be reliably traced to a known cell. These orphan fragments were small (median length 0.38 µm) and rare (4.13±6.05 per dataset). Orphan fragments represent 0.18% of the total neurite length and 0.13% of all synapses and were excluded from analysis.

Modulatory neurons distinguish themselves from non-modulatory neurons by distinct features of their chemical synapses^58^. Chemical synapses come in two varieties: classical synapses, containing mostly clear synaptic vesicles, are made by all non-modulatory neurons; modulatory synapses, containing mostly dense-core vesicles (DCVs), are made by all modulatory neurons.

Classical synapses were identified by a characteristic presynaptic swelling containing a pool of clear vesicles adjacent to a dark presynaptic active zone on the inside of the membrane^29^. Each presynaptic active zone was annotated as the presynaptic partner of one chemical synapse. Cells adjacent to the active zone, within 100nm in xyz dimension, was identified as its potential postsynaptic partners. Annotation included considerations for additional characteristics. Presynaptic swellings were also typically characterized by a small number of DCVs at the periphery of the active zone-associating synaptic vesicle cloud, the presence of mitochondria, as well as the cadherin-like junctions between the pre- and postsynaptic partner cells^59^. Some postsynaptic partners exhibit morphological features such as swelling or postsynaptic densities that resemble the signature PSDs at the mammalian glutamatergic synapses.

Modulatory synapses appear as periodic varicosities along the modulatory neuron’s neurite, each filled with a cloud of DCVs. Some modulatory synapses are devoid of clear synaptic vesicles; some have a small numbers of clear vesicles in these varicosities. Most DCV-specific varicosities did not have presynaptic active zones; the small amount of presynaptic active zones were often not associated with vesicles^58^.

Gap junctions were partially annotated and not subjected to the consensus scoring process due to limitations of current sample preparation protocols^29^.

Final synapse annotations for all datasets are available at http://nemanode.org/. Only chemical synapses with presynaptic active zones were included for subsequent analyses.

### Classification of neuron types

We followed conventional neuronal type classification^14^, with modifications based on structural features revealed in this study and other studies.

Neurons were classified as motor neurons if they primarily made synapses onto muscles. Neurons were classified as sensory if they had specialized sensory processes and/or were previously reported to have sensory capabilities. Neurons were classified as interneurons if most of their connections were to other neurons. Neurons were classified as modulatory if they make chemical synapses that contained mostly large dark vesicles, or, if they had been previously reported to use following neurotransmitters: serotonin (AIM, HSN), dopamine (ADE, CEP), or octopamine (RIC)^60,61^. Some neurons exhibit features corresponding to more than one type. In this case, they were classified based on their most prominent feature. A summary of the classification of each neuron and their justification is provided in Table S1.

### Volumetric reconstruction of cellular processes

We computed the precise shape of every neurite and muscle process in each EM image based on the skeleton tracing that was performed in CATMAID and a machine learning algorithm that recognized cellular boundaries. In brief, the algorithm expanded all skeleton nodes in each section until they fully filled the images of all labelled cells.

Cellular borders were predicted by a shallow Convolutional Neural Network (CNN) that builds on *XNN*^62,63^, a recently developed high performance system which computes convolutions on CPUs, to achieve border prediction throughput of ∼10MB/s^64,65^. Node expansion was computed with a dedicated Cilk-based code^66^ that parallelized the Dijkstra graph search algorithm. Code optimization allowed us to perform node expansion of an entire EM section in memory by a single multithreaded process. Each software thread expanded an individual skeleton. Each pixel is attributed to a given cell by computing a generalized form of distance, taking into account the minimum number of cellular border pixels that must be traversed in a path connecting pixel and node. The generalized distance is computed using graph theory and concurrent data structures.

Volume traces were imported into VAST^67^ for manual proof reading. At least 1,120 person-hours were spent proofreading the volumetric expansions. It was not possible to perform volumetric reconstruction on dataset 7 due to weak membrane contrast.

### Quantification of chemical synapse size

Coordinates of all chemical synapses were exported from CAT-MAID^57^ and imported into VAST^67^ using custom scripts. The presynaptic active zone for every synapse was manually segmented throughout every section where it was visible. The size of monadic synapses is represented by the volume of the presynaptic active zone. At polyadic synapses, we estimated the relative strengths of postsynaptic partners by the proportion of postsynaptic area that each partner occupies at each synapse. We performed a Monte Carlo simulation of neurotransmitter diffusion from the presynaptic active zone, and quantifying the proportion of these particles that reached each potential postsynaptic partner using the three-dimensional geometry of the EM reconstruction. Synapse size for each postsynaptic partner was calculated by multiplying the total volume of the presynaptic active zone by the proportion of particles that hit the membrane of each postsynaptic partner.

### Data processing for analysis

Volumetric neuron traces were exported from VAST^67^ and imported into MATLAB. EM artefacts were manually corrected. To calculate the contact area of each adjacent cell pair, we performed two-dimensional morphological dilation of every traced segment across extracellular space until neighbouring segments made contact within 70 pixels (140-280nm). Expansion was restricted to the edge of the nerve ring. The total contact area was calculated as the sum of adjacent pixels for each segment in all sections. Contacts between cell bodies at the periphery of the neuropil were excluded.

Neuron skeletons and synapses were exported from CAT-MAID using custom Python scripts, and imported into Python or MATLAB environments for analyses. The module detection analysis was performed in MATLAB. Other analyses were implemented with custom Python scripts using SciPy and Statsmodels libraries for statistics. Post-embryonically born neurons were excluded from analyses related to classification of connections, feedforward information flow, and modules.

For analyses related to neurites, both processes of neurons and muscles in the nerve ring were included. The neurite length was calculated using the smoothened skeleton of each neurite. The skeleton was smoothed by a moving average of 10 skeleton nodes after correction of major alignment shifts. Spine-like protrusions were defined as any branch shorter than the 10% of the average neuron length. For analyses related to information flow, separating connections into feedforward, feedback, and recurrent, connections to muscles were excluded since they are all feedforward.

### Classification of connections

A total of 3113 connections (averaging 1292 per dataset) were assigned as stable, variable, or developmentally dynamic. 1647 connections (averaging 323 per dataset) had no more than two synapses in two or more datasets and were left-right asymmetric. These connections were classified as *variable*. The 1466 remaining connections were pooled by left-right cell pairs, resulting in 658 pair connections. The number of synapses in each pair connection was tested for relative increase or decrease across maturation (Spearman’s rank correlation, corrected for multiple comparisons using the Benjamini–Hochberg correction). Pair connections showing a significant change and at least a 5-fold change in synapse number from birth to adulthood were classified as *developmentally dynamic*. When a connection is absent from dataset 1 and 2, but exists in later datasets, we consider it to have increased more than 5-fold (an ‘infinite’ increase). Remaining pair connections were considered *stable* if they were present in at least 7 datasets, and *variable* if present in fewer than 7 datasets. The 5-fold change cutoff is based on the overall expansion in synapse number from early L1 to adulthood. Occasionally, connections were near the cutoff for developmentally dynamic versus variable connections. How they are categorized does not affect any overall pattern in our connectome analysis due to the extremely small number.

### Comparison to the original *C. elegans* adult connectome

The original adult hermaphrodite brain connectome annotated by White et al.^14,44^ was taken from wormatlas.org (dataset N2U). Because individual muscles were not traced in the original annotation, we completed this dataset by tracing through all head muscles using the scanned EM micrographs hosted by wormatlas.org. Individual muscles arms were identified by their characteristic location within the brain, which were confirmed by tracing the arms back to their cell body in several datasets. This augmented dataset (referred to as “N2U, White et al., 1986”) was used for subsequent comparison.

The wormatlas.org hosts a re-annotated version of the wiring of the N2U connectome, which includes synapses to individual muscles from (^24^). We noted errors in muscle identification and synapse annotation in this reannotation. We corrected some errors so that we could perform comparisons with our analysis. Specifically, we corrected the identity of the following muscle pairs VL1-VL2, VR1-VR2, DL2-DL3, DR2-DR3, DL5-DL6, DR5-DR6, VL5-VL6, and VR5-VR6. Other mistakes in tracing and synapse annotation could not be corrected. For example, muscles DL7 and DL8 were not traced at all in the brain, and only one of more than 50 synapses onto muscle VR2 (named as VR1 in Cook et al. 2019) was annotated. This minimally corrected dataset, referred to as “N2U, Cook et al., 2019” was used for subsequent comparison.

For both N2U datasets, we only included neurons and neurites within the same regions used for our datasets for comparison.

### Community structure analysis

Weighted stochastic blockmodeling (WSBM)^32^ was used to define modules individually for all eight connectomes. In this approach, modules are optimized on the likelihood of observing the actual network from the determined modules (log-likelihood score) based on two exponential family distributions. We chose the probability of establishing connections to follow a Bernoulli distribution and the synapse number for each connection to follow an exponential distribution. These distributions fit the data best according to the log-likelihood score and resulted in leftright cell pairs being assigned to the same modules.

In order to find a stable and representative number of modules for each connectome, we used a consensus-based model-fitting approach, similar to previously described^68^. First, to ensure unbiased coverage of the parameter space, we fitted the model independently 300 times using an uninformative prior for each potential number of modules (k = 1, …, 8). This procedure was repeated 100 times to yield a collection of models with concentrated and unimodally distributed log evidence scores. To improve the stability of the models on multiple runs, we increased the parameters for a maximum number of internal iterations to 100. For each dataset, we chose the number of modules whose collection of models had the highest mean posterior log-likelihood score. For dataset 2 the second-highest score was selected, as the number of modules otherwise conflicted with adjacent time points.

Finally, for each dataset, we found a representative consensus module assignment that summarized all 100 models^68^. In brief, considering all 100 models, we calculated the frequency of each cell being assigned to each module, and used this as a new prior to fit another 100 models. This procedure was repeated until convergence, when the consistency of the 100 models was larger than 0.95.

### Statistics

Statistical methods were not used to predetermine sample sizes. Spearman’s rank was used for all correlations (Fig. 2d-2f, S3c, S6d, S6e, S9e and S9f) and time series (Fig. 2g, 3 and S5b-S5d). Two-tailed Z-test was used to compare proportions (Fig. 4c, 5c). To determine if developmentally dynamic connections were over- or underrepresented, the proportions of developmentally dynamic connections between each cell type were compared to the total proportion of developmentally dynamic connections throughout all cell types (Fig. 4c, **??**). Kruskal-Wallis test followed by pairwise Mann-Whitney U tests were used for comparisons of more than two unpaired categories (Fig. 4b, 5b, S6f and S9c, **??**). For figure panels with more than three categories, only categories statistically different from all others were labelled (Fig. 4b, S9c, **??**. For figure panels with multiple comparisons, the reported p-values were FDR adjusted using Benjamini–Hochberg correction.

## Supporting information

Supplementary Info 1 - Neuron types

Supplementary Info 2 - Connectivity matrices

Supplementary Info 3 - Synapses sizes

Supplementary Info 4 - Contact matrices

Supplementary Info 5 - Spine-like protrusions

Supplementary Info 6 - Connection classifications

Supplementary Info 7 - Dataset ages

Supplementary Info 8 - Anatomical inconsistencies

## Data availability

All electron microscopy images and volumetric reconstructions are available at bossdb.org/project/witvliet2020. Connectivity matrices for all datasets are available at www.nemanode.org and as supplementary info.

## Data and code availability

All scripts and files used to generate all figures are available at https://github.com/dwitvliet/nature2021.

## ACKNOWLEDGEMENTS

We thank Valeriya Laskova for assistance in developing EM samples. We thank Bob Harris for assistance with high-pressure freezing. We thank Marianna Neubauer, David Kersen, Anabelle Paulino, Manusnan Suriyalaksh, Amelia Srajer, Maggie Chang, Sean Ihn, and Jade Ho for help with imaging. We thank Albert Cardona, Ignacio Arganda-Carreras and Jenny Qian for guidance on EM alignment. We thank Steven Cook, Christine Rehaluk, and Mona Wang for synapse annotation in some datasets. We thank Jade Ho, Christopher Morii-Sciolla, Isis So, Min Wu and Chi Yip Ho for help with generating ground truth and proofreading for volumetric reconstruction. We thank Alexander Matveev, Lu Mi, and Hayk Saribekyan for help with generating and applying algorithms used in volumetric reconstruction. We thank Jerry Wang and Danqian Cao for help with statistical analyses. We thank Albert Lin, Chris Tabone, and Vivek Venkatachalam for setting up and supporting the server for tracing and annotation. We thank Soomin Maeng and Dylan Fong for assistance with the development of www.nemanode.org. We thank Mona Wang, Wesley Hung, and Jun Meng for examining iBLinC and NAFT methods and the labs of the Hannes Buelow and David Miller for sharing reagents and discussions. We thank members of the Zhen, Samuel, and Lichtman labs for comments. We especially thank Guangwei Si and Lav Varsney for critical reading and suggestions. We thank David Hall, Jagan Srinivasan, and Albert Cardona for advice in early phase of this project.

J.K.M. was supported by National Science Foundation Physics of Living Systems (NSF 1806818). B.M. was supported by the Mount Sinai Foundation. J.W.L. was supported by the National Institute of Mental Health, Silvio Conte Center (P50 MH094271), the National Institutes of Health (U24 NS109102-01), and the Multidisciplinary University Research Initiative (GG0008784). J.W.L., A.D.T.S., and M.Z. were supported by the Human Frontier Science Program (RGP0051/2014). A.D.T.S and M.Z. were supported by the National Institutes of Health (R01-NS082525-01A1). A.D.T.S. was supported by National Institutes of Health Brain Initiative (1U01NS111697-01) and National Science Foundation BRAIN EAGER (IOS-1452593). M.Z. was supported by Canadian Institutes of Health Research (MOP-123250 and Foundation Scheme 154274), the Radcliffe Institute for Advanced Studies, and the Mount Sinai Foundation.

## AUTHOR CONTRIBUTIONS

J.W.L, A.D.T.S., and M.Z. conceived and guided the study. Y.M., R.P., and N.S. designed the algorithm for automated volumetric reconstruction ( for correspondence). D.R.B. and R.L.S. designed the pipeline for automated EM acquisition ( for correspondence). Y.W. designed software for EM alignment ( for correspondence). B.M., D.H. and M.Z. generated EM samples. D.W., B.M., J.K.M., D.H., R.L.S, and M.Z. imaged most of the electron micrographs. D.W., B.M., and J.K.M. performed most annotation. D.W. designed and performed most analysis. D.R.B., W.X.K., and Y.L. performed additional experiments and analysis. A.D.C. guided early cell identification and annotation. D.W., J.W.L, A.D.T.S., and M.Z. wrote the manuscript. All authors reviewed the manuscript.

## Supplemental figures

**Video 1. Fly-through of an adult EM dataset**.

**Video 2. Volumetric reconstruction of an adult dataset**.

**Video 3. Individual neurons across maturation**.

**Video 4. Modules in the adult brain**.

**Table S1.** Cell types in the nerve ring. This table lists cell type using each neuron, muscle, and glia that contributed to chemical synapses included in our analyses. Each is assigned a cell type using the described criteria (see Methods). We performed volumetric reconstructions of all listed neuron and muscle processes within our datasets. We did not perform volumetric reconstructions of the much thinner glia processes, which our algorithms (see Methods) were unable to reconstruct automatically. Volumetric reconstruction of the 6 GLR glia (cells with a mesodermal origin that may affect neuron-muscle communication) and the 4 CEPsh glia (the sheath cells of the cephalic sensilla that have a neuronal/epidermal origin) will require thinner EM sectioning^69^.

| Class | Members | Type | Integration into nerve ring |
| --- | --- | --- | --- |
| ADA | 2 | inter | embryonic |
| ADE | 2 | modulatory | embryonic |
| ADF | 2 | sensory | embryonic |
| ADL | 2 | sensory | embryonic |
| AFD | 2 | sensory | embryonic |
| AIA | 2 | inter | embryonic |
| AIB | 2 | inter | embryonic |
| AIM | 2 | modulatory | embryonic |
| AIN | 2 | inter | embryonic |
| AIY | 2 | inter | embryonic |
| AIZ | 2 | inter | embryonic |
| ALA | 1 | modulatory | embryonic |
| ALM | 2 | sensory | embryonic |
| ALN | 2 | sensory | post-embryonic |
| AQR | 1 | sensory | post-embryonic |
| ASE | 2 | sensory | embryonic |
| ASG | 2 | sensory | embryonic |
| ASH | 2 | sensory | embryonic |
| ASI | 2 | sensory | embryonic |
| ASJ | 2 | sensory | embryonic |
| ASK | 2 | sensory | embryonic |
| AUA | 2 | sensory | embryonic |
| AVA | 2 | inter | embryonic |
| AVB | 2 | inter | embryonic |
| AVD | 2 | inter | embryonic |
| AVE | 2 | inter | embryonic |
| AVF | 2 | modulatory | post-embryonic |
| AVH | 2 | modulatory | embryonic |
| AVJ | 2 | modulatory | embryonic |
| AVK | 2 | modulatory | embryonic |
| AVL | 1 | modulatory | embryonic |
| AVM | 1 | sensory | post-embryonic |
| AWA | 2 | sensory | embryonic |
| AWB | 2 | sensory | embryonic |
| AWC | 2 | sensory | embryonic |
| BAG | 2 | sensory | embryonic |
| BDU | 2 | inter | embryonic |
| BWM01 | 4 | muscle | embryonic |
| BWM02 | 4 | muscle | embryonic |
| BWM03 | 4 | muscle | embryonic |
| BWM04 | 4 | muscle | embryonic |
| BWM05 | 4 | muscle | embryonic |
| BWM06 | 4 | muscle | embryonic |
| BWM07 | 4 | muscle | embryonic |
| BWM08 | 4 | muscle | embryonic |
| CEP | 4 | modulatory | embryonic |
| CEPsh | 4 | glia | embryonic |
| DVA | 1 | modulatory | embryonic |
| DVC | 1 | inter | embryonic |
| FLP | 2 | sensory | embryonic |
| GLR | 6 | glia | embryonic |
| HSN | 2 | modulatory | post-embryonic |
| IL1 | 6 | motor | embryonic |
| IL2 | 6 | sensory | embryonic |
| OLL | 2 | sensory | embryonic |
| OLQ | 4 | sensory | embryonic |
| PLN | 2 | sensory | post-embryonic |
| PVC | 2 | inter | embryonic |
| PVN | 2 | modulatory | post-embryonic |
| PVP | 2 | inter | embryonic |
| PVQ | 2 | modulatory | embryonic |
| PVR | 1 | inter | embryonic |
| PVT | 1 | inter | embryonic |
| RIA | 2 | inter | embryonic |
| RIB | 2 | inter | embryonic |
| RIC | 2 | modulatory | embryonic |
| RID | 1 | modulatory | embryonic |
| RIF | 2 | inter | embryonic |
| RIG | 2 | inter | embryonic |
| RIH | 1 | inter | embryonic |
| RIM | 2 | inter | embryonic |
| RIP | 2 | inter | embryonic |
| RIR | 1 | inter | embryonic |
| RIS | 1 | modulatory | embryonic |
| RIV | 2 | motor | embryonic |
| RMD | 6 | motor | embryonic |
| RME | 4 | motor | embryonic |
| RMF | 2 | motor | post-embryonic |
| RMG | 2 | modulatory | embryonic |
| RMH | 2 | motor | post-embryonic |
| SAA | 4 | sensory | embryonic |
| SDQ | 2 | sensory | post-embryonic |
| SIA | 4 | motor | embryonic |
| SIB | 4 | motor | embryonic |
| SMB | 4 | motor | embryonic |
| SMD | 4 | motor | embryonic |
| URA | 4 | motor | embryonic |
| URB | 2 | sensory | embryonic |
| URX | 2 | sensory | embryonic |
| URY | 4 | sensory | embryonic |

**Table S2.**
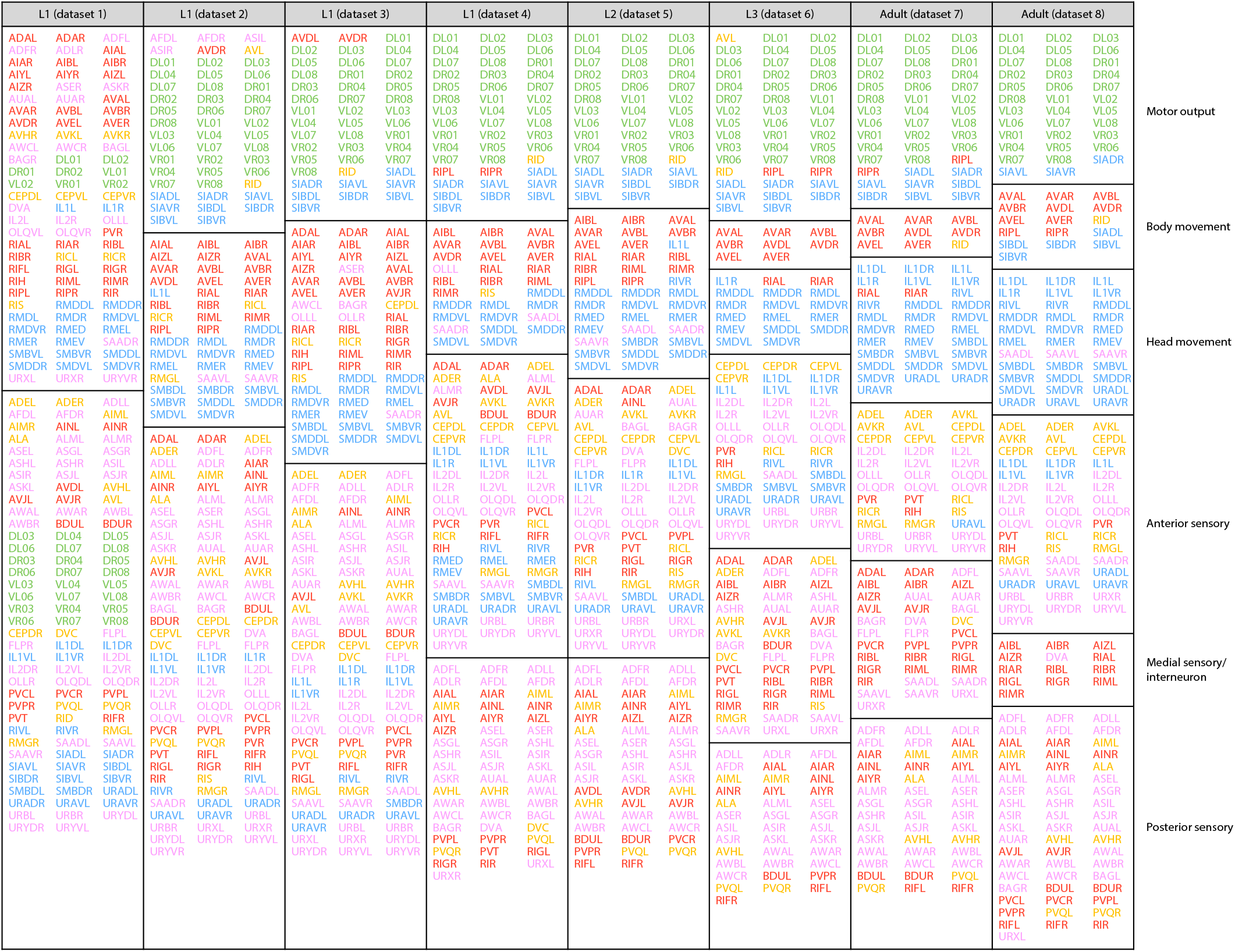
Members of modules detected by WSBM colored by type.

**Table S3.** Optimal number of modules detected by WSBM using subsets of connections.

| **Connections included** | **Dataset 1 (L1)** | **Dataset 2 (L1)** | **Dataset 3 (L1)** | **Dataset 4 (L1)** | **Dataset 5 (L2)** | **Dataset 6 (L3)** | **Dataset 7 (Adult)** | **Dataset 8 (Adult)** |
| --- | --- | --- | --- | --- | --- | --- | --- | --- |
| All connections | 2 | 3 | 3 | 4 | 4 | 6 | 6 | 6 |
| Non-variable connections | 2 | 2 | 2 | 2 | 2 | 4 | 5 | 5 |
| Stable connections | 2 | 2 | 2 | 2 | 2 | 2 | 2 | 2 |

**Figure S1.**
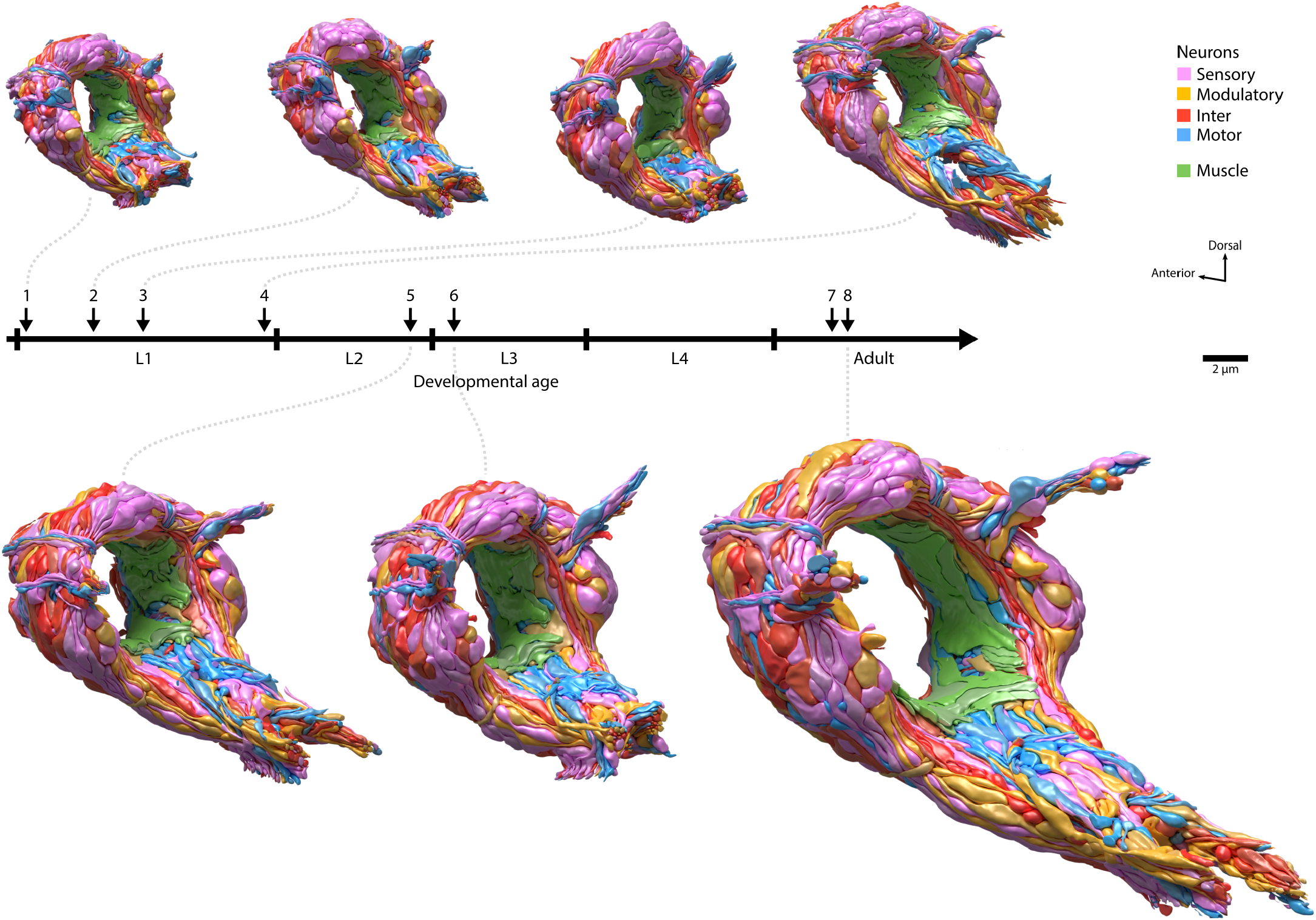
Volumetric models for seven *C. elegans* brains at respective developmental stages. All models include the complete neuropil and muscle fibers of the brain, consisting of the nerve ring and ventral ganglion. Glia processes are not included. Cells are colored by type.

**Figure S2.**
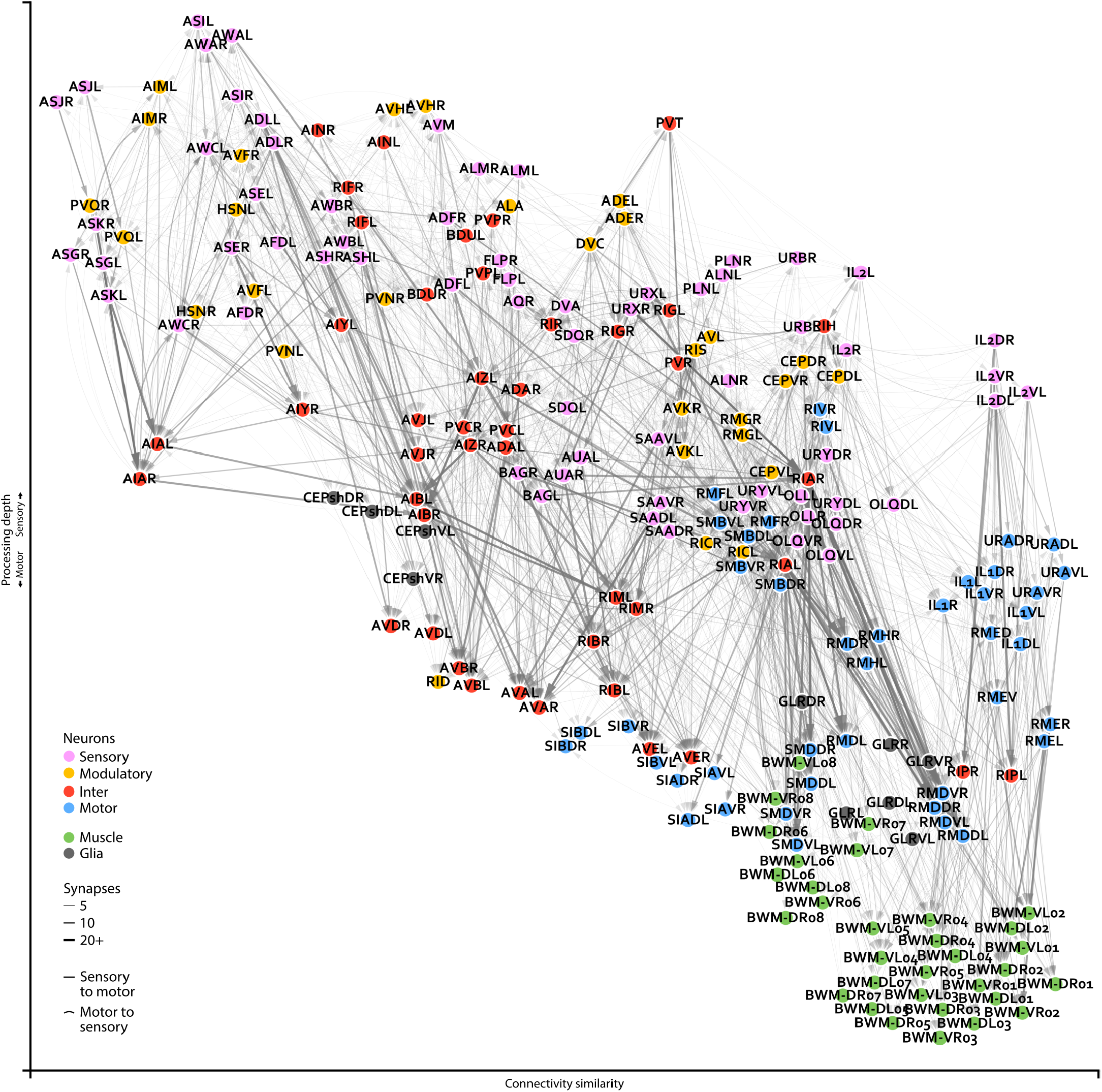
Closeup of an adult brain connectome. Wiring diagrams for an adult connectome (dataset 8). Each circle represents a cell. Circle colour denotes cell type. Each line represents a connection with at least one chemical synapse between two cells. Line width indicates synapse number. Straight lines direct information from sensory to muscle layers whereas curved lines direct information in reverse. Cell coordinates are represented as in Fig. 1b, with overlapping cells manually separated.

**Figure S3.**
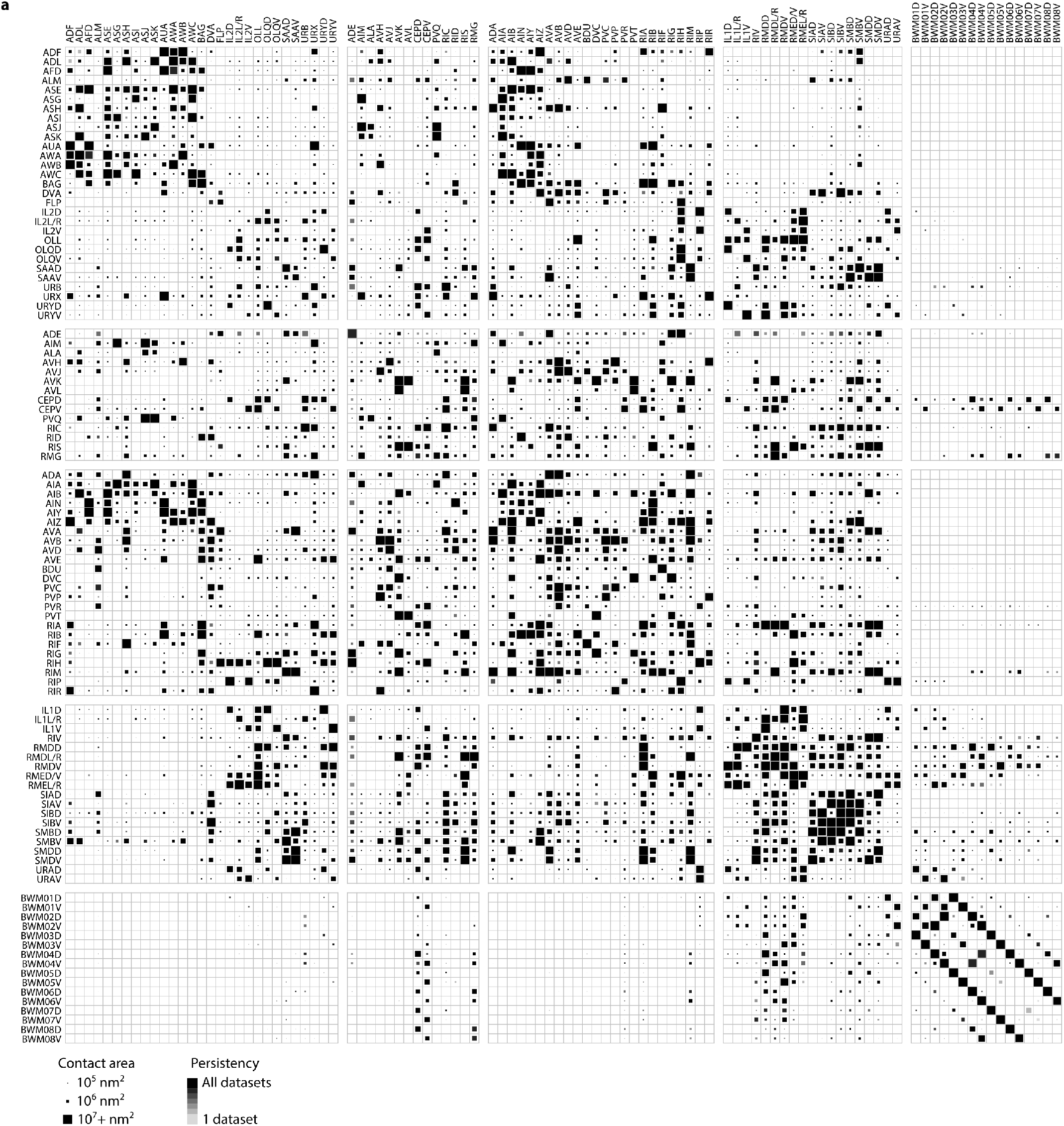
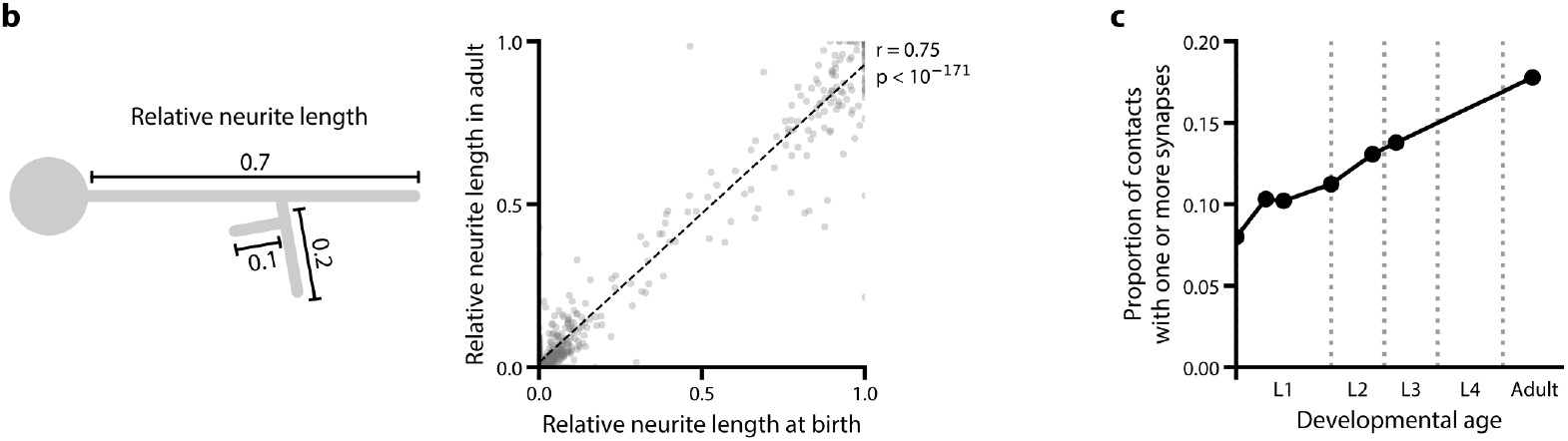
A physical contact matrix between neurites and muscle fibers in seven volumetrically reconstructed *C. elegans* brains. **a**. Cells are pooled by left-right pairs. The physical contact size is represented by the largest value from the seven datasets. **Neurites grow while maintaining overall brain geometry. b**. Correlation of the relative neurite length of each branch between L1 (dataset 1) and adult (dataset 8). The length of each neurite is normalized against the total neurite length of the neuron. p = 9.4×10^−172^, r = 0.75, n = 947, Spearman’s rank correlation. **c**. Proportion of physical contacts in the brain that harbors at least one chemical synapse at respective developmental time points.

**Figure S4.**
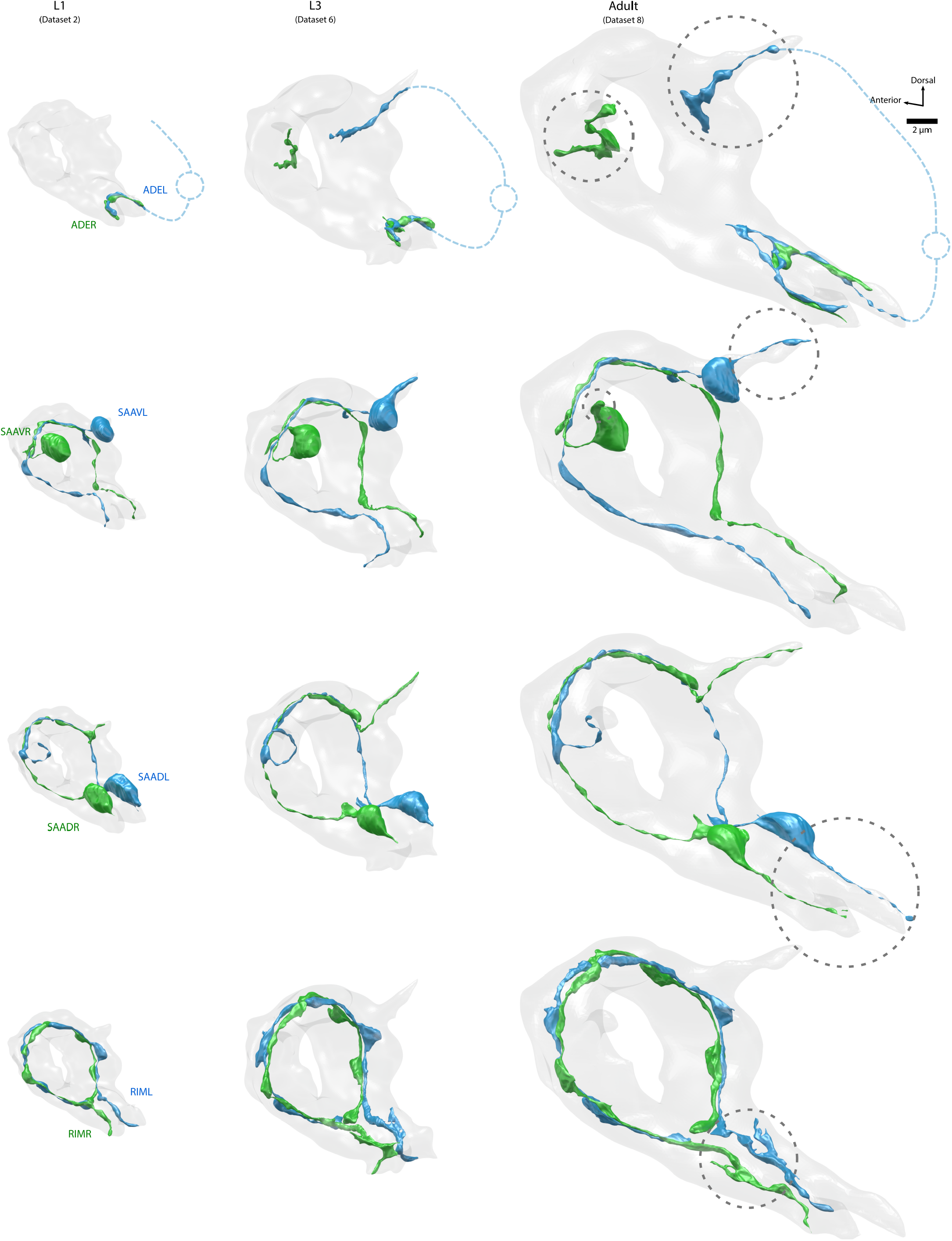
Three neuron classes grow new neurites after birth. Volumetric models of ADE, SAAV, SAAD, and RIM in L1 (dataset 2), L3 (dataset 6), and adult (dataset 8). These neurons pairs grow new major branches, highlighted by dotted gray circles. ADE’s new branches sprout outside the brain; regions not volumetrically reconstructed are denoted by a dotted blue line.

**Figure S5.**
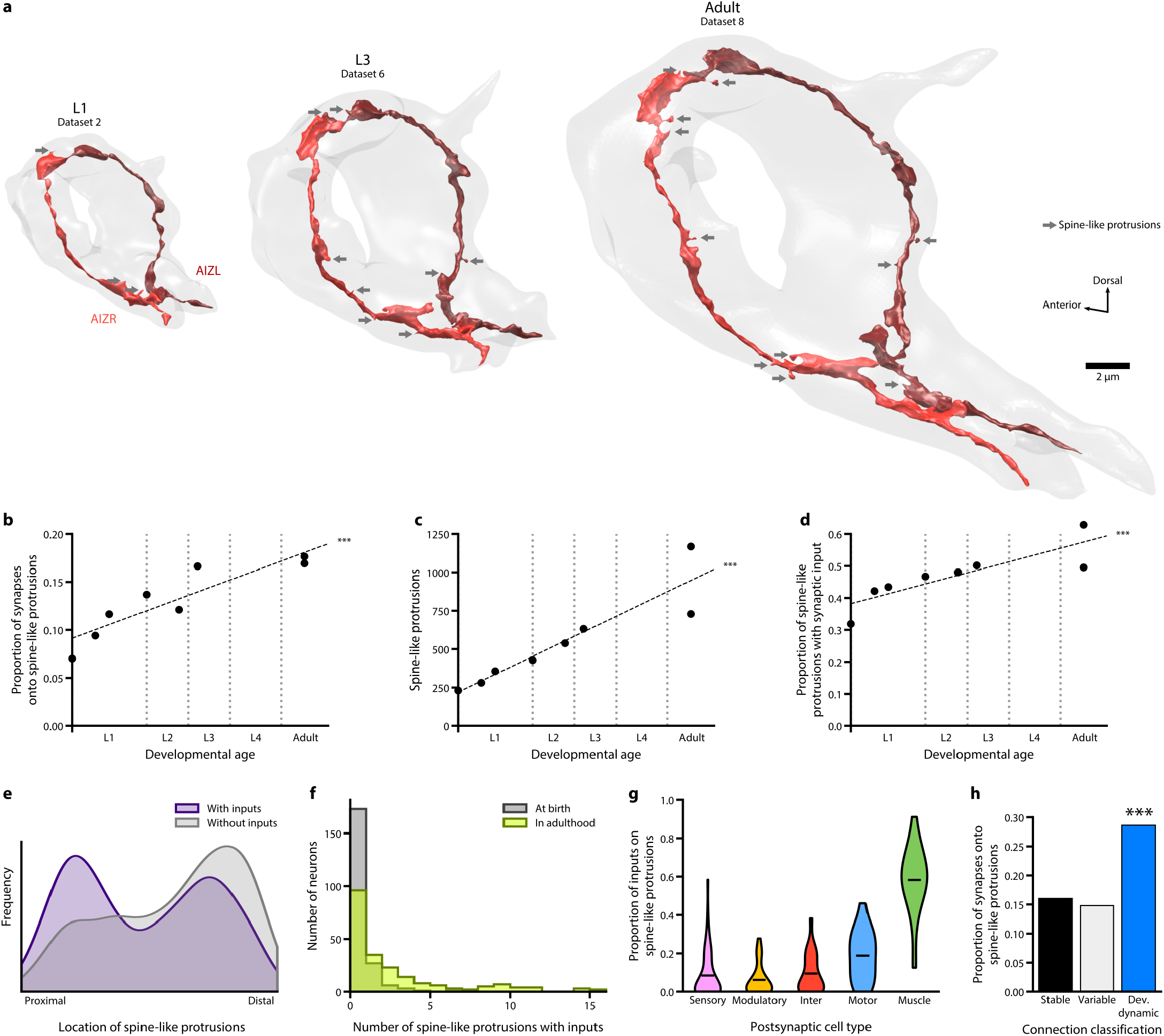
Prevalence, location, and synaptic distribution of spine-like protrusions. **a**. 3D reconstructions of one neuron class (AIZL and AIZR) across maturation. The overall geometry was maintained, whereas the number of spine-like protrusions (grey arrows) increased over time. **b**. Proportion of postsynaptic spine-like protrusions increases across maturation. *** p = 6.5×10^−5^, Spearman’s rank correlation. **c**. Total number of spine-like protrusions in the brain increases across maturation. *** p = 5.3×10^−7^, Spearman’s rank correlation. **d**. Proportion of synapses with at least one spine-like protrusion postsynaptic partner increases across maturation. *** p = 1.8×10^−4^, Spearman’s rank correlation. **e**. Distribution of spine-like protrusions by location, with the entry of the neurite into the brain as the most proximal, and the exit or terminal end of the neurite the most distal. **f**. Number of spine-like protrusions that oppose a presynaptic terminal per neuron at birth (averaged between datasets 1 and 2) and in adulthood (averaged between datasets 7 and 8). **g**. Proportion of presynaptic inputs onto spine-like protrusions per neuron in adulthood (averaged between datasets 7 and 8), grouped by their cell type.

**Figure S6.**
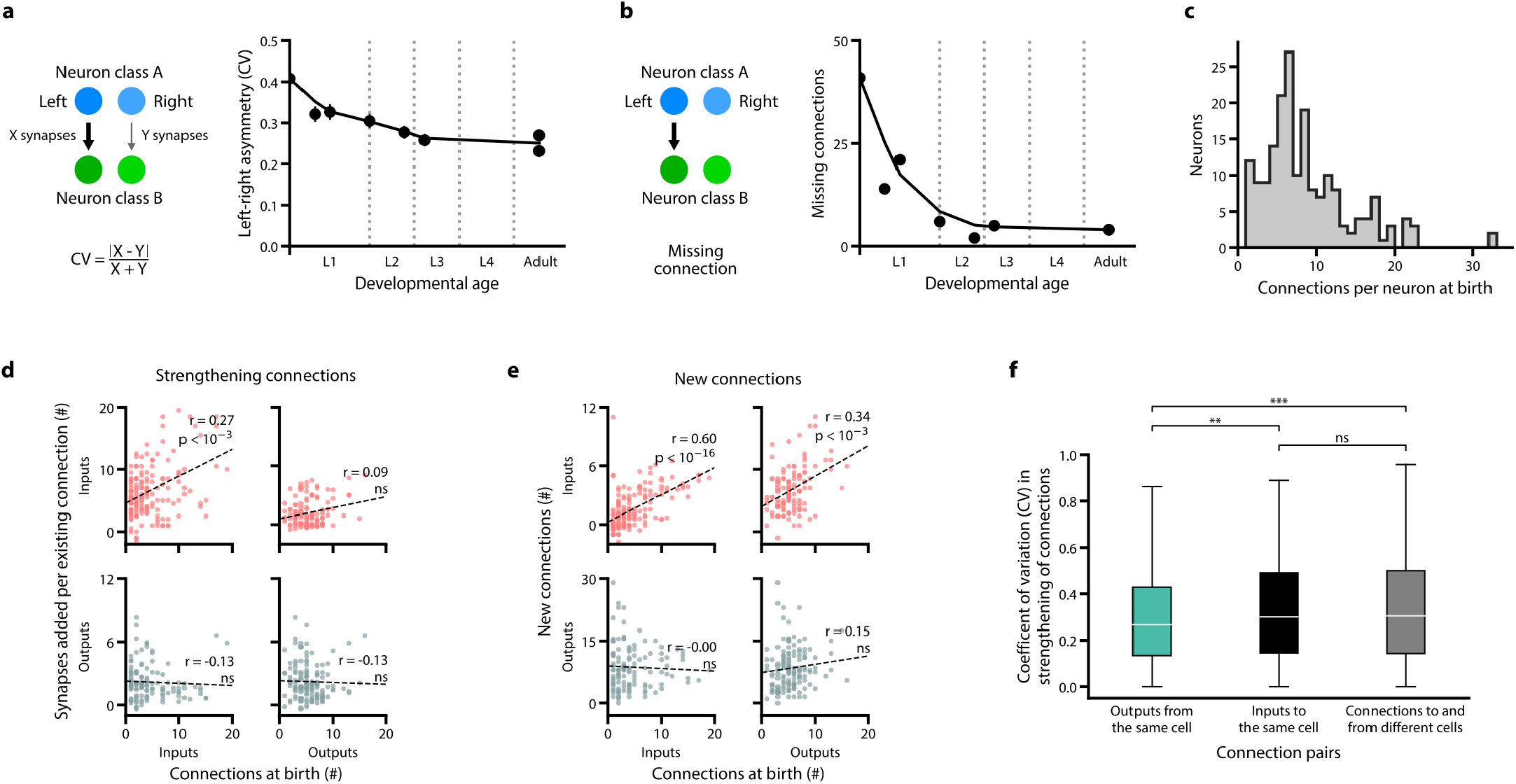
Most connectivity asymmetry at birth is eliminated during L1. **a**. Connectivity asymmetry decreases from birth to adulthood, most significantly during L1. Asymmetry is defined as the coefficient of variation (CV) in synapse number between left-right cell pairs. Error bars indicate SE. **b**. Total number of missing connections decreases from birth to adulthood, most significantly during L1. One connection refers to a cell making at least one chemical synapse to another cell. A missing connection is defined as a connection absent in only one dataset and from one side of the brain. **Non-uniform distribution of connections and strengthening of connections across maturation c**. Distribution of the total number of input and output connections per neuron at birth. Some neurons have more connections than others.**d**. Upper panels: neurons with more input connections at birth are more likely to strengthen these connections during maturation. Left: the number of input connections at birth (dataset 1) is positively correlated with their synapse number increase by adulthood (averaged between datasets 7 and 8). p = 1.6×10^−17^, n = 166 by the Spearman’s rank correlation. Right: the number of output connections at birth does not predict the synapse number increase at input connections by adulthood. p = 0.32, n = 120 by the Spearman’s rank correlation. Lower panels: Neither input connection (left) nor output connection (right) at birth predicts the synapse number increase at output connections by adulthood. left: p = 0.16, n = 120; right: p = 0.12, n = 141 by the Spearman’s rank correlation. Each point represents one cell. **e**. Upper panels: neurons with higher number of input connections (left) or output connections (right) at birth (dataset 1) are more likely to establish new input connections by adulthood (averaged between datasets 7 and 8). Left: p = 5.4×10^−4^, n = 166; right: p = 1.7×10^−4^, n = 120 by the Spearman’s rank correlation. Lower panels: Neither the input (left) or output (right) connection number at birth predicts the likelihood to establish new output connections by adulthood. Left: p = 1.00, n = 120; right: p = 0.08, n = 141 by the Spearman’s rank correlation. Each data point represents one cell.**f**. The relative number of synapses added to existing connections is correlated between outputs of the same cell compared to connections to and from different cells. The relative number of synapses added is quantified as the fold increase of synapse number from birth (dataset 1) to adulthood (averaged between datasets 7 and 8). ns (not significant) p = 0.24, ** p = 2.3×10^−3^, *** p = 2.5×10^−5^, Mann–Whitney U test, FDR adjusted using Benjamini–Hochberg correction (n_outputs_ = 753, n_inputs_ = 1203, n_other_ = 90709). Center line, median; box limits, upper and lower quartiles; whiskers, 1.5x interquartile range; outliers not shown.

**Figure S7.**
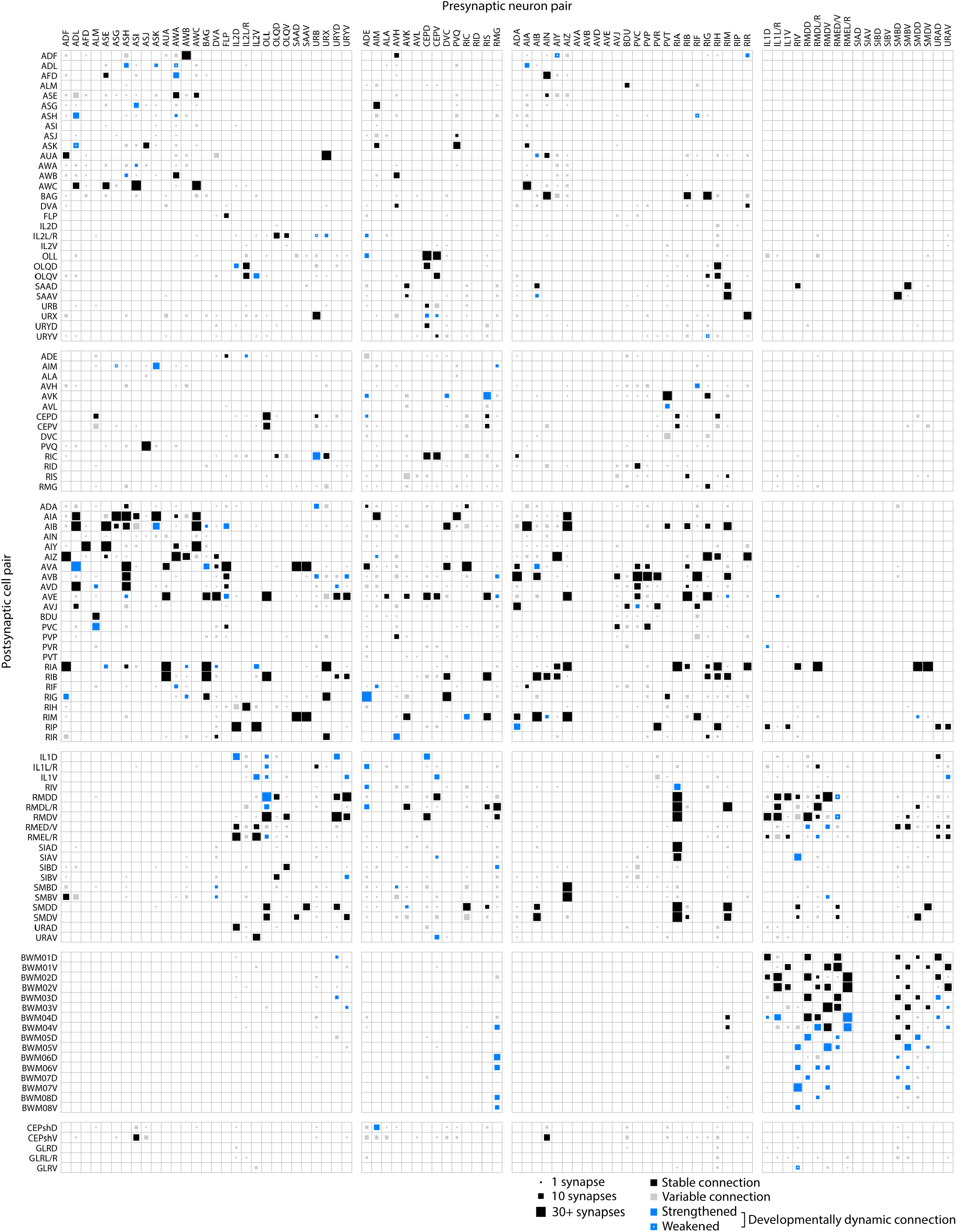
Connectivity matrix of the *C. elegans* brain throughout maturation. A connectivity matrix that includes all connections observed in eight *C. elegans* brains. Cells are pooled by left-right pairs. The size of each connection represents its largest synapse number in any dataset. Stable, developmentally dynamic, and variable connections are colour-coded by their classification (see Methods).

**Figure S8.**
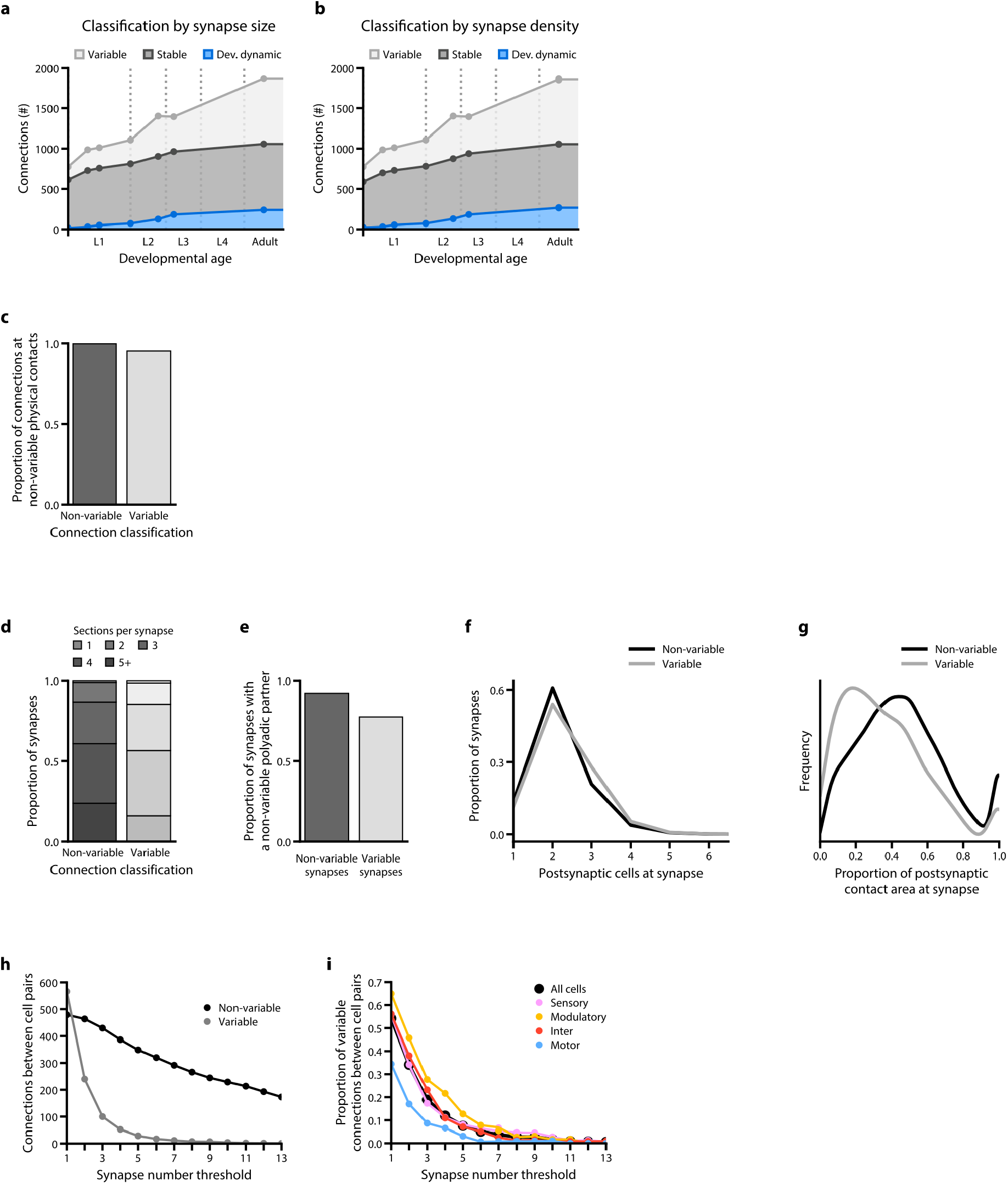
A connectome has prevalent variable connections. **a**. Composition of stable, developmentally dynamic, and variable connections in each dataset classified by synapse size. **b**. Composition of stable, developmentally dynamic, and variable connections in each dataset classified by synapse density, defined by the total synapse number divided by the cable length of the input neuron). **Prevalence of variable connections is not caused by over-annotation of ambiguous synapses c**. High proportions of both variable and non-variable (stable and developmentally dynamic) connections form at non-variable physical contacts. A physical contact is defined as variable when it is absent from more than one of the seven datasets. **d**. Synapses that constitute non-variable and variable connections, sorted by EM section numbers that the presynaptic active zone encompasses. All synapses in seven volumetrically segmented datasets are included. Synapses comprising variable connections are marginally smaller that those comprising non-variable connections, but no threshold can be set to remove exclusively the variable connections. **e**. Proportion of synapses that form a polyadic synapse with synapses of the stable connections. A marginally smaller portion of synapses that comprise variable connections (78%) than those comprising non-variable connections (93%) reside in this configuration. Therefore, variable connections are fortuitous accidents of synapse annotation. **f**. Synapses comprising non-variable and variable connections sorted by the number of post-synaptic partners. They exhibit similar distribution from being monoadic to polyadic. Non-variable connections have marginally more polyadic synapses than variable connections (20% vs 28% for dyadic, and 61% vs 54% for triadic synapses, respectively). No threshold by postsynaptic partner number can be set to filter variable connections. **g**. Proportion of postsynaptic contact area occupied by each postsynaptic partner at each synapse. Synapses comprising variable connections on average occupy less of the postsynaptic area than synapses comprising non-variable connections, but no threshold can be set to only exclude variable connections. **All threshold removes both variable and non-variable connections. h**. Total number of non-variable (stable and developmentally dynamic) and variable connections in adulthood (averaged between datasets 7 and 8) upon thresholding by different synapse numbers. No synapse number provides a filter for specific removal of variable connections: all removes both variable and stable connections. **i**. Thresholding connections by synapse number leaves substantial proportion of variable connections for all cell types. Non-uniform distribution of variable connections remains when connections with low synapse numbers are removed.

**Figure S9.**
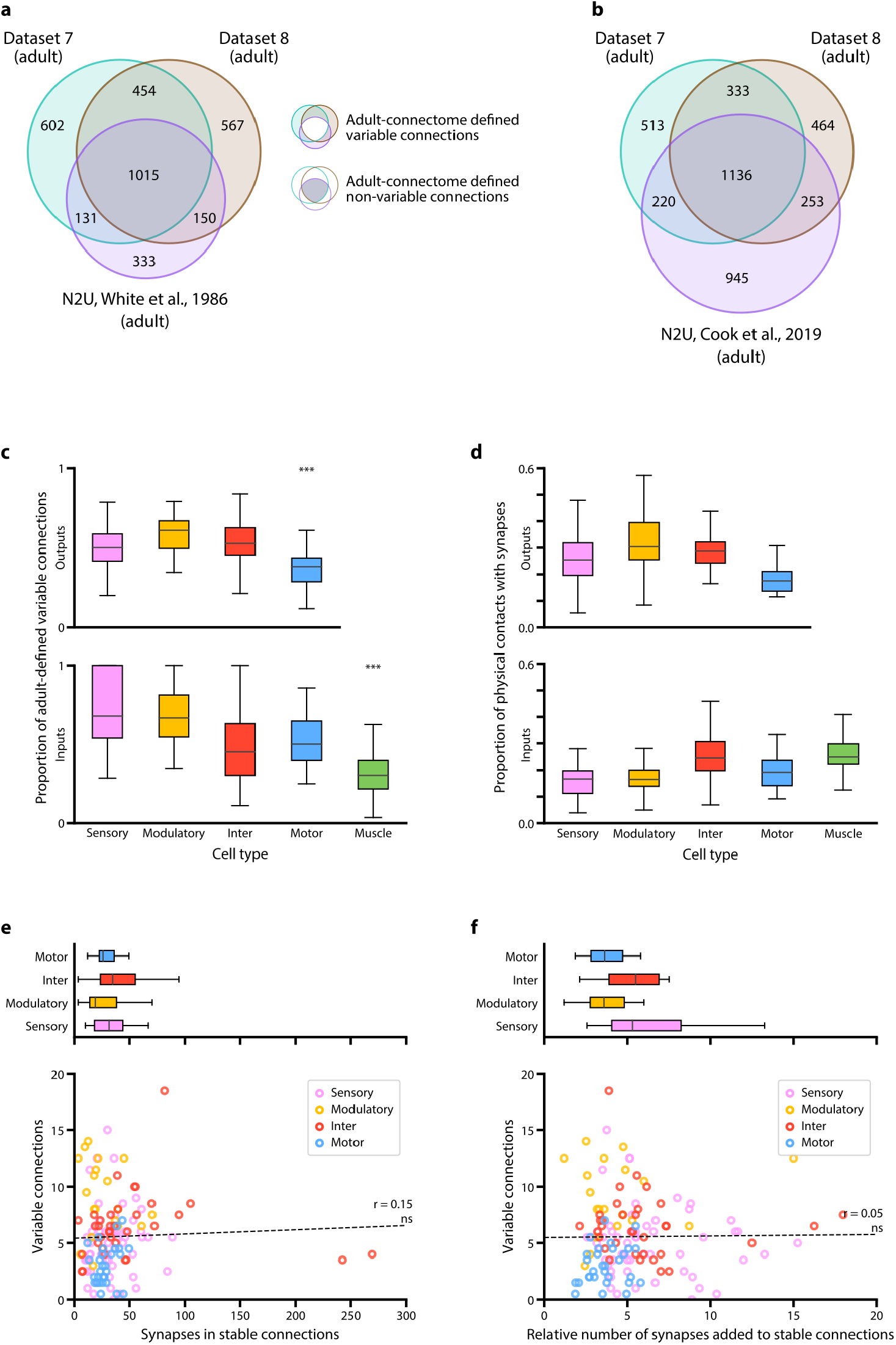
Comparison of multiple adult connectomes reveals extensive variability in connectivity. **a** Shared and unique connections for three adult connectomes: dataset 7, dataset 8, and N2U **(a)** annotated by White et al.^14^, illustrated in the Venn diagram. Connections of all synapse numbers are included for comparison (Methods). **Re-annotation of N2U increased its variability. (b)** Re-annotated of the N2U adult connectome (Cook et al.^24^)) added 1109 new connections that disproportionally enlarged its pool of unique connections (see Methods). Only 16% contributed to connections shared by three connectomes. This may imply the application of different annotation criteria from the original annotation. **Propensity of forming variable connections correlates with cell type. c**. Comparison between the proportion of adult connectome-defined variable and non-variable connections for each cell type. Adult-defined non-variable connections include the connections that are present in both of our adult datasets as well as the original connectome annotated by White et al.^14^. Cell types with significantly higher or lower proportions of variable connections are denoted, ** p < 10^−2^, *** p < 10^−3^, n = 28-65, Mann–Whitney U test, FDR adjusted using Benjamini–Hochberg correction. Center line, median; box limits, upper and lower quartiles; whiskers, 1.5x interquartile range; outliers not shown. **d**. The low variability of connections from motor neurons to muscles cannot be simply explained by saturation of their physical contacts by synapses. Physical contacts are not saturated for connections for any cell type. Motor neurons, which have the lowest proportion of variable connections (Fig. 4b), are not restricted by few available potential synaptic partners. Center line, median; box limits, upper and lower quartiles; whiskers, 1.5x interquartile range; outliers not shown. **e-f**. Higher variability for certain cell types could also not be simply explained by a fixed probability of an erroneous connection by neurons that exhibit abundant synapse formation. **e**. Top: The number of synapses for stable output connections by cell types. Modulatory neurons, which exhibit a higher proportion of variable connections than other cell types (Fig. 4b), do not exhibit more synapses per stable connection. Center line, median; box limits, upper and lower quartiles; whiskers, 1.5x interquartile range; outliers not shown. Bottom: The number of variable connections formed by a cell does not correlate with the strength of its stable output connections. Each data point represents one cell. ns (not significant) p = 0.08, r = 0.15, n = 139, Spearman’s rank correlation coefficient. **f**. Top: The relative number of synapses added to existing stable output connections by cell types. Connections from modulatory neurons, which have a higher proportion of variable connections than other cell types (Fig. 4b), do not exhibit higher increase in synapse number than connections from other cell types. Center line, median; box limits, upper and lower quartiles; whiskers, 1.5x interquartile range; outliers not shown. Bottom: The number of variable connections formed by a cell does not correlate with the number of synapses added to existing stable output connections from birth to adulthood. The relative number of synapses added is quantified as the fold increase of synapse number from birth (dataset 1) to adulthood (averaged between datasets 7 and 8). Each data point represents one cell. ns (not significant) p = 0.56, r = 0.05, n = 139, Spearman’s rank correlation coefficient. For panels d-f, the synapse number for the adult brain (averaged between datasets 7 and 8) is shown.

**Figure S10.**
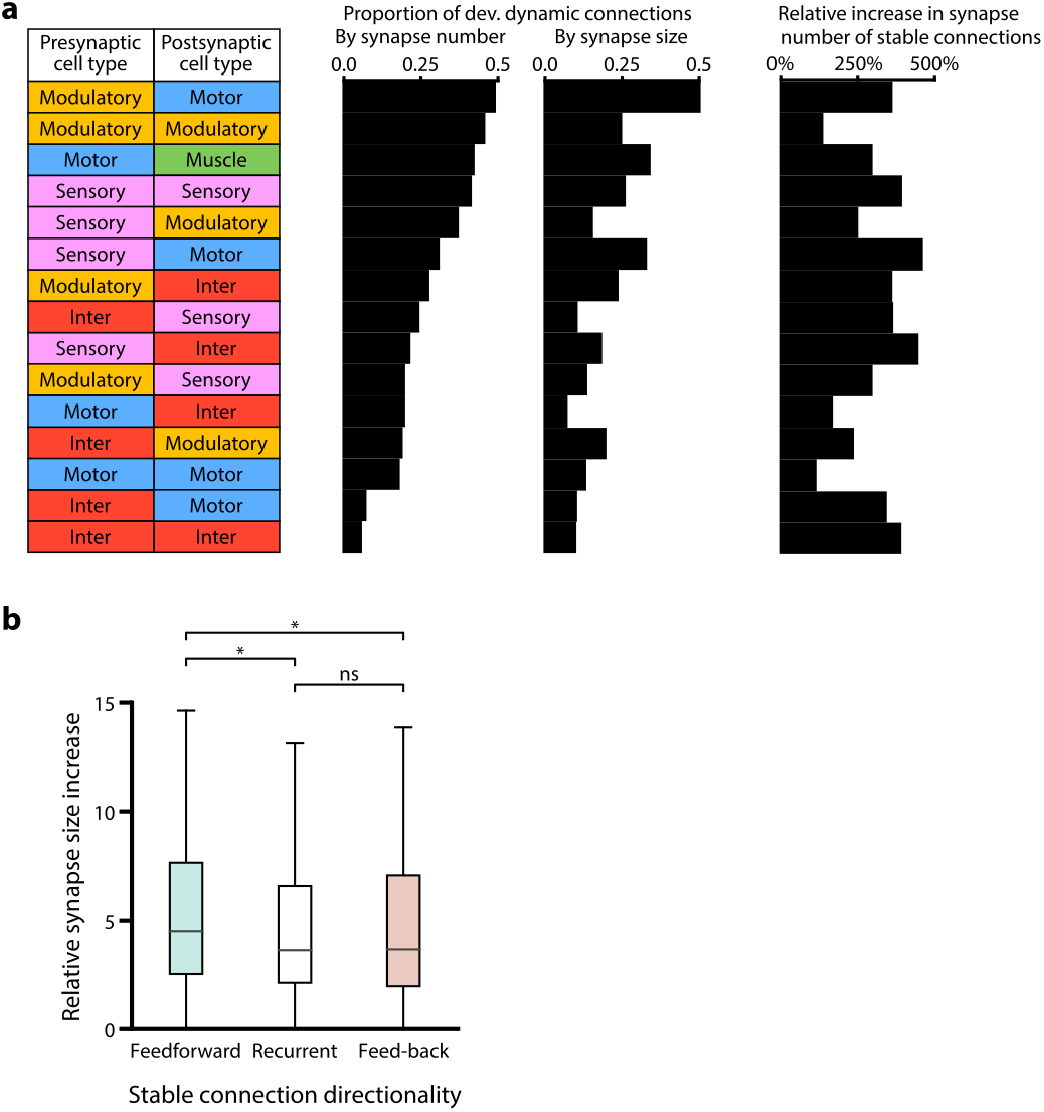
Stability of interneuron connections and strengthening of feedforward connections are revealed by assessing connection strength by synapse size. **a**. Proportion of developmentally dynamic connections by cell type, when connection strength changes were evaluated by either synapse number (left) or synapse size (middle). Connections between interneurons are the most stable regardless of how synapse weight was evaluated. Right panel: Developmental stability of connections is not correlated with the extend of synapse number increase from birth (averaged between datasets 1 and 2) to adulthood (averaged between datasets 7 and 8). **Spine-like protrusions are significantly enriched at developmentally dynamic connections. b**. Proportion of synapses with spine-like protrusions that comprise stable, variable, and developmentally dynamic connections. Developmentally dynamic connections have the highest proportion. *** p < 10^−24^, two-tailed Z-test, FDR adjusted using Benjamini–Hochberg correction (n_stable_ = 10059, n_variable_ = 2169, n_dev. dynamic_ = 1611). **c**. Fold increase of summed synapse size for stable connections from birth (averaged between datasets 1 and 2) to adulthood (averaged between datasets 7 and 8). Feedforward connections are strengthened more than feedback and recurrent connections. ns (not significant) p = 0.39, * p < 0.05, Mann–Whitney U test, FDR adjusted using Benjamini–Hochberg correction (n_feedforward_ = 301, n_recurrent_ = 229, n_feedback_ = 107). Center line, median; box limits, upper and lower quartiles; whiskers, 1.5x interquartile range; outliers not shown.

**Figure S11.**
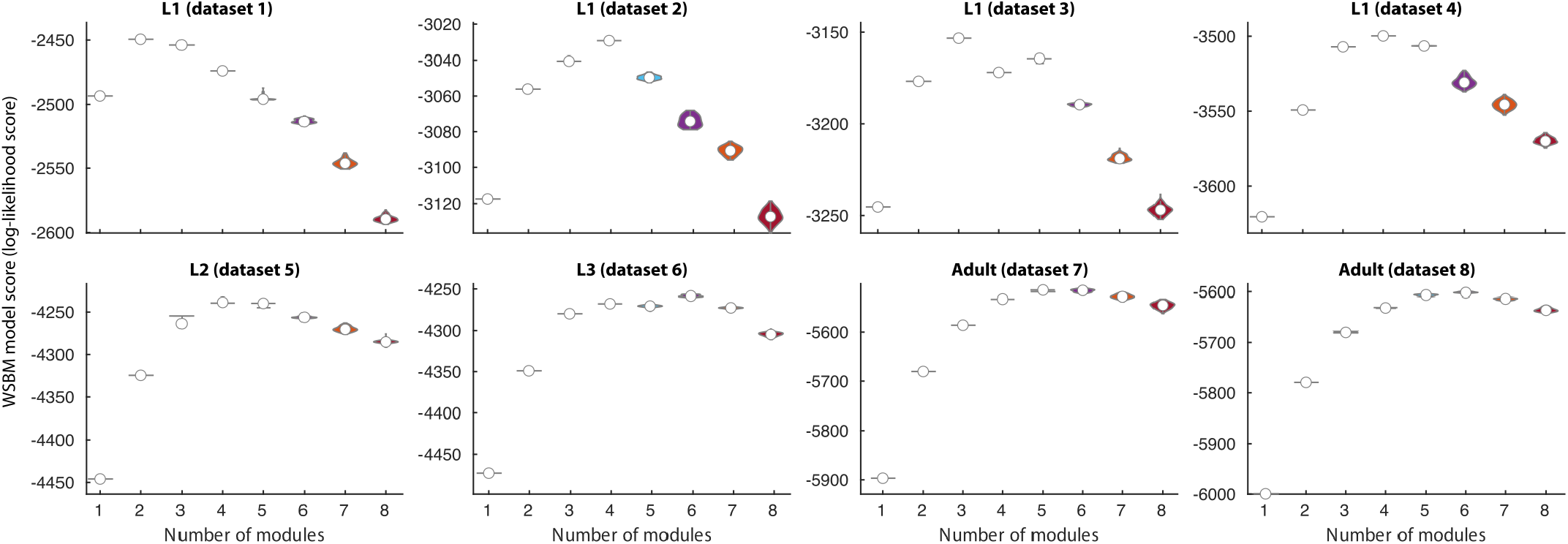
Cell modules across maturation. **a**. The log-likelihood score for each WSBM model (see Methods). **b**. The deviation between the connectome and each synthetic network generated from the best WSBM model, measured by the mean KS energy (see Methods). A lower deviation indicates a better match between the actual connectome and network generated from the model. Adult datasets show a clear preference to more than 5 modules, while juvenile datasets do not.

