## Supplementary Info 7 - Dataset ages for "Connectomes across development reveal principles of brain maturation"

### Estimation of the developmental age of datasets

The developmental age of each sample was established based on the described temporal cell division pattern exhibited by wild-type (N2) larva raised at 25 degrees C (Sulston and Horvitz 1977).

Dataset 1: L1 at birth. No Q cell division, which occurs ~3 hours post-hatching (hrs) after birth. Q cell nuclei are symmetrical, before nuclei migration, which occurs ~2 hrs after birth, placing this sample so close to birth, estimated to be ~0 hrs.

Dataset 2: L1 estimated to be 5 hrs. Q cell divided but no H1 division, which occurs at~7.5 hrs after birth. We found that PVC neurites were partially grown out and SAA posterior neurites had not begun to grow at this time point, indicating an age slightly more than halfway between dataset 1 and 3.

Dataset 3: L1 at 8 hrs. With H1 just completing its division and P5/6 starting their migration, this sample was placed at ~8 hrs after birth, when both events take place.

Dataset 4: L1 at the very end of the larval stage, near the end of the L1 lethargus. It is estimated to be 16hrs. It has two layers of cuticle. Both P11.aaa and P12.aaa have divided. V5R.p is in the midst of division, and H1.a has not yet divided; all happen at ~16 hrs.

Dataset 5: L2 towards the end of the larval stage (23hr). SML/R have not divided, which occurs at ~29 hrs. It has 40 gonad cells, and a slight double cuticle that indicates the end of L2. However, its gut lumen contains food, placing this sample shortly before entering L2 lethargus, which occurs at ~23 hrs.

Dataset 6: L3 at 27 hrs. Based on the partial outgrowth of the RMF neurites, which is born at 23 hrs, this sample was estimated to be ~27 hrs.

Datasets 7 and 8: Young adults at 45hr. Both samples have adult cuticles but are relatively small compared to other adults. The exact ages of the two young adult samples are uncertain, so they are treated as equals for analyses.
