## Supplementary Info 8 - Anatomical inconsistencies for "Connectomes across development reveal principles of brain maturation"

### Anatomical inconsistencies between samples

A few anatomic inconsistencies are observed in some datasets, likely due to heterogeneity or imprecision of development processes. These events do not have an impact on overall connectivity, as all non-variable connections between individual neuron classes were conserved. Neurons in these datasets did not exhibit more variable connections either.

Dataset 2: CEPDL cell body is shifted to the anterior ganglion.

Dataset 3: RIFL neurite terminates prematurely laterally, not reaching the dorsal midline.

Dataset 4: RIH cell body is shifted to the anterior ganglion. PVCL and PVCR neurites both go right-handedly around the nerve ring, appearing as PVCR.

Dataset 5: RMHL and RMHR neurites both transverse right-handedly around the nerve ring, appearing as RMHR. ADAL terminates prematurely at a dorsal sub-lateral position, not reaching the dorsal midline.

Dataset 6: PVR neurite is fragmented.

Dataset 7: RIFL and RIFR neurites both transverse right-handedly around the nerve ring, appearing as RIFR.
